# Neurocomputational evidence that conflicting prosocial motives guide distributive justice

**DOI:** 10.1101/2022.05.02.490266

**Authors:** Yue Li, Jie Hu, Christian C. Ruff, Xiaolin Zhou

## Abstract

In the history of humanity, most conflicts within and between societies have originated from perceived inequality in resource distribution. How humans achieve and maintain distributive justice has therefore been an intensely studied issue. However, most research on the corresponding psychological processes has focused on inequality aversion and has been largely agnostic of other motives that may either align or oppose this behavioral tendency. Here we provide behavioral, computational, and neuroimaging evidence that distribution decisions are guided by three distinct motives - inequality aversion, harm aversion, and rank reversal aversion – that interact and in fact can also deter individuals from pursuing equality. At the neural level, we show that these three motives are encoded by separate neural systems, compete for representation in various brain areas processing equality and harm signals, and are integrated in the striatum, which functions as a crucial hub for translating the motives to behavior. Our findings provide a comprehensive framework for understanding the cognitive and biological processes by which multiple prosocial motives are coordinated in the brain to guide redistribution behaviors. This framework enhances our understanding of the brain mechanisms underlying equality-related behavior, suggests possible neural origins of individual differences in social preferences, and provides a new pathway to understand the cognitive and neural basis of clinical disorders with impaired social functions.

## Introduction

Throughout the course of human history, from Aristotle’s *Nicomachean ethics* to *Marxism* and the *Declaration of Independence*, the pursuit of fairness and equality has been a cornerstone of social justice and has led to the continuing development of human societies (Binmore, 1998, 2005; Fehr and Rockenbach, 2004). Fairness principles not only affect everyone’s daily life (e.g., salary distribution at the workplace) but also shape social political ideology and therefore social welfare (e.g., tax policies and health-resource distribution policies) (Baron, 1995; Offer and Pinker, 2017). In line with this universal importance, people usually approach issues of distributive justice from the perspective of fairness norms (Rawls, 1999) which are often considered as the most fundamental principles by which humans distribute resources (Sanfey et al., 2003; Hsu et al., 2008; Tricomi et al., 2010). This view is increasingly supported by empirical evidence that people not only help disadvantaged parties to gain more equally distributed outcomes (Lowery et al., 2009; Hu et al., 2015; Xie et al., 2017) but also punish behaviors that violate the fairness norm (Masclet and Villeval, 2008; Corradi-Dell’Acqua et al., 2013; Ginther et al., 2016; Jordan et al., 2016; Tan and Xiao, 2018; Lo Gerfo et al., 2019; House et al., 2020).

However, fairness norms and inequality aversion alone cannot fully account for choices in situations requiring resource redistribution, which can in principle also reflect very different motives (Hsu et al., 2008). For example, imagine that two colleagues have made the same contributions to a project, but their employer decided to give one of them 1,000 dollars as bonus and the other one only 100 dollars (i.e., A: $1000 / B: $100). Most people would feel frustrated by such an unequal distribution (Sanfey et al., 2003; Xiao and Houser, 2005; Corradi-Dell’Acqua et al., 2013) and would generally be willing to help the disadvantaged (Lowery et al., 2009; Tricomi et al., 2010; Yu et al., 2014b), albeit within certain limits. For example, most people would be happy to transfer 200 dollars from the advantaged to the disadvantaged to achieve a more equal distribution (i.e., A: $800 / B: $300), but many people would be reluctant to transfer 700 dollars since this would reverse the initial dis/advantage rankings of each party (i.e., A: $300 / B: $800). This gives an example of the core motive conflicts associated with distributive justice, which in real life are known to lead to intense debates, for example on how to increase taxation on wealthy people and organizations while at the same time protecting everyone’s interests and maintaining social order (Iosifidi and Mylonidis, 2017). This real-life example emphasizes the necessity to explore the boundaries of inequality aversion and to understand the natural limits of what people would do in the name of “fairness” (Lévy-Garboua et al., 2008; Dimick et al., 2016; Argentiero et al., 2021).

In situations like the above dilemma, and taxation debates in general, a primary aim is to reduce social inequality. However, this always involves trade-offs between inequality aversion (which favours transfer from the wealthy to the less well-off) and at least two other motives that support the status quo - **harm aversion** (Baron, 1995; Crockett et al., 2014) and **rank reversal aversion** (Xie et al., 2017). That is, moral decision studies suggest that people generally do take into account the ‘do-no-harm’ principle and have a tendency to avoid helping one group at the expense of harming another group, even when the benefits outweigh the harm (Baron, 1993, 1995; Crockett et al., 2014). This entails that people are reluctant to redistribute wealth by transferring money from the advantaged party to the disadvantaged party (Leliveld et al., 2008; Wu et al., 2012). Supporting this tendency, people are averse to overturn stable hierarchies in a society even though such pre-existing hierarchies conflict with their inequality aversion (Fernandez and Rodrik, 2004; Magee and Galinsky, 2008; Zitek and Tiedens, 2012). During wealth redistribution, it is widely observed that people are indeed anchored to the initially unequal distribution and support such inequality to avoid reversal of pre-existing income rankings (Xie et al., 2017). Thus, while harm aversion and rank reversal aversion can also be seen as prosocial motives (in that they promote social welfare), they can work against inequality aversion and deter people from pursuing more equal distributions.

To establish the boundaries of these different motives (i.e., inequality aversion, harm aversion, and rank reversal aversion), we have to uncouple them and examine how each of them contributes to redistribution behaviors in situations where they are in conflict. However, previous studies most often employed paradigms specialized to study each motive in isolation, potentially biasing participants to act in line with just one factor. For instance, since in most of previous paradigms, the participant either played as a victim of unfair distribution (De Quervain et al., 2004; Tricomi et al., 2010; Yu et al., 2014b) or played as an irrelevant third-party to punish intentional norm violations (Strobel et al., 2011; Hu et al., 2015), motives to maximize one’s own interests or to punish norm violators to maintain social justice may have amplified individuals’ inequality aversion in these situations. Moreover, due to the limitations of previous paradigms and the narrow focus of extant econometric models (Fehr and Schmidt, 1999; Charness and Rabin, 2002), it is difficult to differentiate harm aversion and rank reversal aversion from inequality aversion, and to clarify how humans weigh between these motives to make redistribution decisions. The trade-off between these multiple prosocial motives may actually challenge the basic assumption of econometric social preference models that distribution behaviors depend on ultimate outcomes rather than the changes between ultimate and initial outcomes (Fehr and Schmidt, 1999; Charness and Rabin, 2002).

In the current study, we aim to develop an integrated approach to examine how these different prosocial motives (i.e., inequality aversion, harm aversion, and rank reversal aversion) interact with each other to guide wealth redistribution choices. Specifically, we present a novel paradigm and an unbiased modelling approach that allows us to establish the boundaries and relative strengths of each motive, and to elucidate the neural mechanisms underlying their effects on redistribution. That is, we employ functional magnetic resonance imaging (fMRI) to clarify how information relevant for the different motives (e.g., inequality and harm signals) is represented and integrated in the human brain when people make redistribution decisions. One hypothesis emerging from the literature is that equality-related information may be represented in the reward system (e.g., striatum and vmPFC, Cappelen et al., 2014; De Quervain et al., 2004; Strobel et al., 2011; Tricomi et al., 2010; Yan Wu et al., 2014) and that individuals’ preferences related to equality-seeking can be predicted by this activity and the connectivity strengths between these regions and other systems (e.g., prefrontal regions and anterior insula) (Yu et al., 2014b; Glass et al., 2015; Hu et al., 2015). With respect to harm aversion and rank reversal aversion, the literature suggests that social-cognition (e.g., temporal parietal junction (TPJ)) and executive control systems (e.g., prefrontal regions) may underlie expression of these motives, since these structures have been found to be associated with greater preferences to minimize others’ loss or pain (i.e., harm aversion or guilt aversion) (Chang et al., 2011; Crockett et al., 2017). Moreover, stimulation-induced activation of DLPFC can causally increase individuals’ guilt feeling to harm others and substantially dampen harm brought upon others during wealth distribution (Nihonsugi et al., 2015). Thus, TPJ and DLPFC may be sensitive to information concerning harm to others, which may be expressed as harm aversion and rank reversal aversion in the current paradigm.

After identifying the systems involved in representing the information relevant for each motive, we then examined how these motives are weighed and coordinated in the brain to guide redistribution decisions. To answer this question, we focus on how neural systems responding to different signals interact with each other to affect decisions in line with the latent motives. This allows us to differentiate between two potential scenarios regarding the motive-weighing process. On the one hand, while similar neural responses to equality signals have been observed in the striatum across different contexts, the connectivity of striatum with other brain regions has varied (Tricomi et al., 2010; Yu et al., 2014b). Therefore, one possible scenario is that equality signals are represented invariantly in the human brain, but conveyed differentially to other systems, or weighed less heavily for choice, during conflicts with other motives (Scenario 1: Conflict gating of equality signals). On the other hand, previous studies have suggested that neural sensitivity to equality signals can depend on how strongly individuals weigh equality under different contexts, and that equality signals may in fact only be expressed when individuals’ decisions are actually guided by equality (Gao et al., 2018). Therefore, neural equality representations may actually be suppressed when other motives conflict with inequality aversion (Scenario 2: Conflict inhibition of equality signals).

To address these questions, we developed a novel redistribution game that allowed us to measure individuals’ inequality aversion, harm aversion, and rank reversal aversion during wealth redistribution choices. In the redistribution game, the participant played as a third-party to redistribute wealth between two anonymous strangers. They were first presented with a monetary distribution offer between two strangers (e.g., Initial offer: Person A: Ұ15, Person B: Ұ3) and were told that these initial endowments were allocated randomly by a computer. They could choose between two alternative offers to reach a more equal distribution. Critically, we included two conditions: In the baseline condition, the two alternative offers were both more equal than the initial offer but maintained the payoff ranking across the initially advantaged and disadvantaged person (e.g., Offer 1: Person A: Ұ14, Person B: Ұ4; Offer 2: Person A: Ұ10, Person B: Ұ8). In the critical condition, by contrast, participants were presented with the same initial offer and the same more unequal alternative offer (e.g., Offer 1: Person A: Ұ14, Person B: Ұ4), but with a different alternative offer (e.g., Offer 2: Person A: Ұ8, Person B: Ұ10) that had the same inequality level as the alternative in the baseline condition but reversed the initial relative rankings (Figure 1A & B). If redistribution decisions are only driven by inequality aversion, people would be more likely to choose the more equal offer regardless of whether or not the more equal offer will reverse the initially relative rankings. But if harm aversion and rank reversal aversion are at play, people will choose the more equal offer less often in the critical condition than in the baseline condition. This allows us to capture harm aversion (via participants’ decision weights on how much money is taken away from the advantaged party) and rank reversal aversion (by a binary weight on whether the initial rankings are reversed). Crucially, we set up the offer matrix carefully to make sure our paradigm and model can effectively capture these different motives (for details, please see Materials & Methods section). Combining this experimental design with computational modelling and fMRI approaches, we could distinguish the effect of each motive on redistribution behaviors and the underlying neural mechanisms.

**Figure 1.**
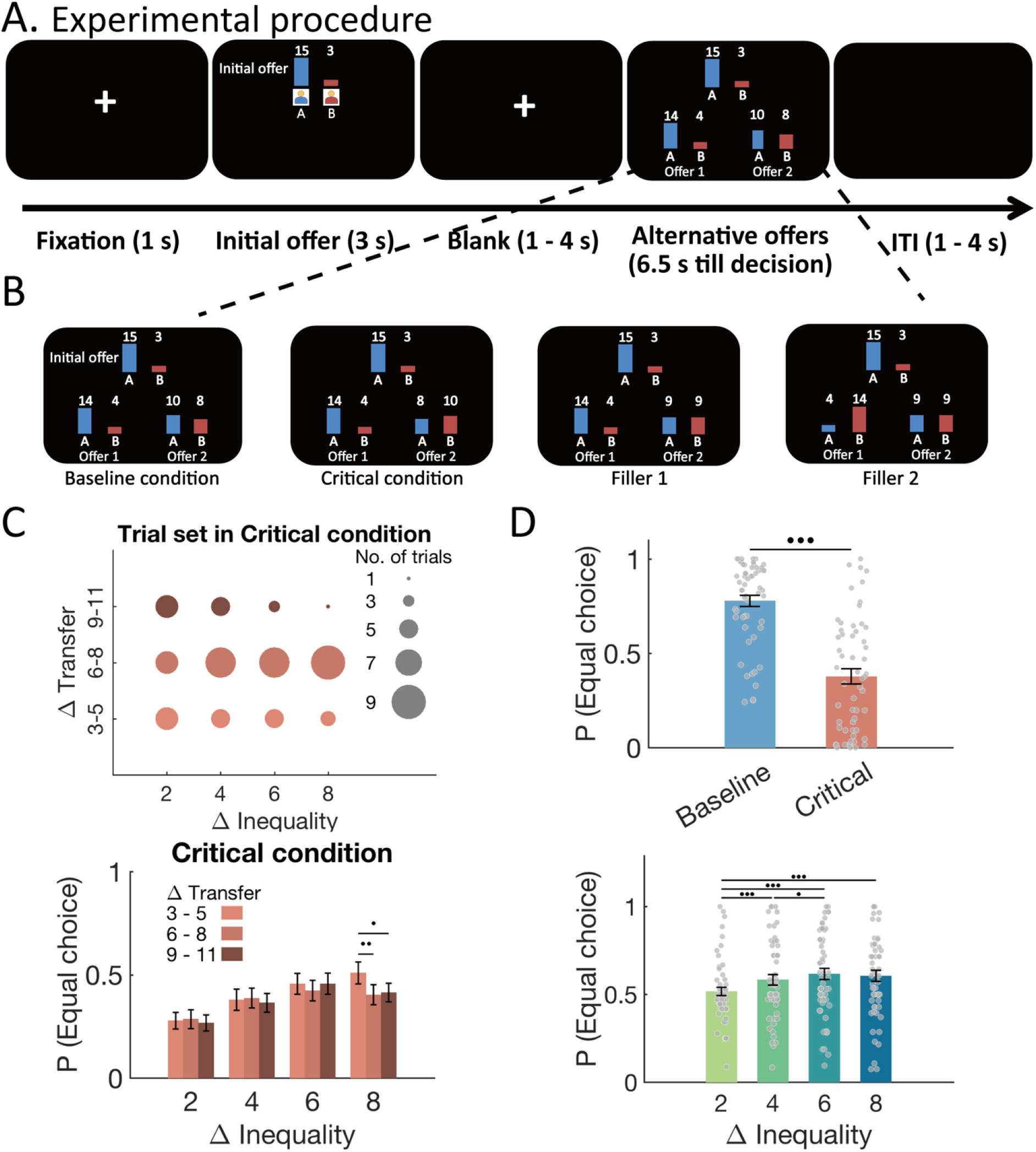
Experimental design and behavioral results. **(A)** The procedure of the task. In each trial, participants were first presented with an unequal monetary distribution scheme (i.e., initial offer) between two anonymous strangers. Then, they could re-distribute the initial scheme by choosing between two alternative offers. Each trial began with a cross fixation on the screen for 1 s. Then, the pictures of the two anonymous strangers together with their initial endowments were presented for 3 s. After a blank screen jittering from 1 to 4 s, the two alternative offers were presented. Participants had to choose one out of the two alternative offers within 6.5 s. After a blank screen jittering from 1 to 4 s, the next trial began. (**B)** Experimental design. In all the trials, both alternative offers were more equally distributed than the initial offer. In the baseline condition (leftmost panel), the initially advantaged party (the person who had more) will still gain more than the initially disadvantaged party (the person who had less) from both alternative offers. In the critical condition (middle left panel), the initially advantaged party would still gain more from the relatively more unequal alternative offer, but would gain less from the relatively more equal alternative offer than the initially disadvantaged party. The inequality levels for both alternative offers were matched across the baseline and critical condition. We also included two types of filler trials in which the more equal alternative offers are always equally divided between the two parties, but the more unequal alternative offer would keep the initially relative rankings in one type of filler trials (middle right panel) and reverse it in the other type of filler trials (rightmost panel). **(C)** To differentiate the effect of inequality and the amount of transferred money (i.e., harm to the advantaged party), we orthogonalized the differences in inequality and the transferred money between the two alternative offers in the critical condition. Top panel, x axis represents the difference in inequality level (Δ Inequality) between the two offers, and y axis represents the difference in transferred money (Δ Transfer) between the two offers. The size of the circle is proportional to the number of trials in each type of Δ Inequality-Δ Transfer combination. Bottom panel, behavioral results in the critical condition, probability to choose more equal alternative offer [P (Equal choice)] is depicted as a function of Δ Inequality and Δ Transfer. The interaction suggested that P (Equal choice) would decrease with the increase of transferred money especially when the inequality difference was high (i.e., 8). For this post-hoc analysis, trials in the critical condition were divided for each level of Δ Inequality into low (i.e., 3-5), middle (i.e., 6-8), and high levels (i.e., 9-11) of Δ Transfer (please note that in the baseline condition, Δ Inequality cannot be differentiated from Δ Transfer). **(D)** Main effects of condition (top panel) and inequality difference (bottom panel) on probability to choose more equal alternative offer. P (Equal choice) significantly decreased when the more equal offer would reverse initially relative ranking (Critical condition) compared with when there was no reverse (Baseline condition), and P (Equal choice) would increase when the more equal offer could reduce the inequality level more strongly. Each dot represents one participant, and error bars represent SEMs. •••, p < 0.001; ••, p < 0.01; •, p < 0.05.

## Results

### Model-free results

To examine whether individuals’ choices reflected only inequality aversion or also the other motives, we first performed a generalized mixed-effect regression of redistribution choices on inequality differences between the two offers (most relevant for inequality aversion), amount taken from the initially more advantaged player in the alternative offer (most relevant for harm aversion), and a binary variable indicating whether the alternative option would reverse the rank between both players (most relevant for rank reversal aversion) (see Materials & Methods for model details). This showed that participants’ redistribution decisions indeed depended on all these three factors: As expected, and indicative of inequality aversion, participants chose the more equal offer more frequently when it more strongly reduced the inequality level (main effect of Δ Inequality with ORE = 1.58, 95% CI [1.37 - 1.83], *p* < 0.001, Figure 1D bottom panel, Table S2) and when the initial inequality was greater (main effect of Δ Initial endowment with ORE = 1.12, 95% CI [1.01 – 1.24], *p* = 0.04, Figure S2A, Table S3). However, individuals’ probability to choose the more equal alternative offer was lower in the critical condition than in the baseline condition (ORE (odds ratio estimate) = 0.37, 95% CI [0.33 - 0.42], P_Baseline_ (Equal choice) = 0.78 ±0.03, P_Critical_ (Equal choice) = 0.38 ±0.04, *t*(56) = 8.88, *p* < 0.001, Figure 1D top panel), demonstrating that rank reversal aversion and harm aversion influence choices independently from inequality considerations (which were matched across the two conditions). Importantly, participants chose the more equal offer less frequently when it entailed larger transfers of money from the advantaged to the disadvantaged party (main effect of Δ Transfer with ORE = 0.46, 95% CI [0.43 – 0.50], *p* < 0.001, Figure S2B, Table S4), highlighting the effect of harm aversion on top of rank reversal aversion. This was also clearly evident in a two-way interaction between Δ Inequality and Δ Transfer (ORE = 0.69, 95% CI [0.50 – 0.96], *p* = 0.03), and a three-way interaction between Δ Inequality, Δ Transfer, and condition (ORE = 1.44, 95% CI [1.16 – 1.79], *p* < 0.001).

To visualize and more directly compare the patterns of the effects in the big regression model, we divided all trials based on condition, Δ Inequality, and Δ Transfer, and inspected how individuals’ choices varied as functions of these variables. Since we had orthogonalized the differences in initial endowment and in transfer/inequality between the two alternative offers, the effects reported here are not confounded by the effect of initial endowment (for details in experimental design, please see Figure 1C top panel and Figure S1). These post-hoc tests confirmed that harm aversion indeed had a stronger effect on redistribution for higher levels of inequality difference (i.e. Δ Inequality = 8, *t*_low vs middle, Δ Inequality = 8_ (56) = 2.71, *p*_low vs middle, Δ Inequality = 8_ = .009, *t*_low vs high, Δ Inequality = 8_ (56) 2.36, *p*_low vs high, Δ Inequality = 8_ = .022, Figure 1C bottom panel, Table S5). In addition, we also observed a significant interaction between Δ Initial Endowment and Δ Transfer (ORE = 1.19, 95% CI [1.10 – 1.28], *p* < 0.001), suggesting that when the inequality level of the initial payoff is low, harm aversion leads people to choose the more equal offer less often when this requires greater monetary transfers and thus more strongly harms the initially advantaged person (Figure S2C, Table S6).

### Model-based results

To better understand the effects and interactions of the different motives on redistribution behaviors, we developed, fitted, and compared four families of computational models identifying how people weigh the different motives to make redistributive decisions. We focused these analyses on the critical condition, which in contrast to the baseline condition allowed us to differentiate inequality aversion from harm aversion and rank reversal aversion (pleased see supplementary methods for rationale and Materials & Methods section for technical details of model selection and estimation).

The control model M1 only considered inequality aversion, whereas M2-M4 considered combinations of inequality aversion and the other motives. The simplest model M1 followed the classical inequality aversion model proposed by Fehr and Schmidt (1999) in which people assign values to the outcomes of all parties but devalue the inequality they experience for any kinds of distribution. Since both parties of the distribution are anonymous strangers for the participant, we considered the absolute value of the payoff difference between the two parties as the inequality level of the two offers:

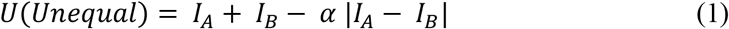

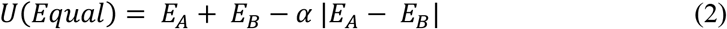

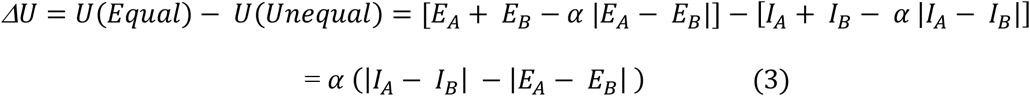

where *I_A_*(*I_B_*) is the payoff of the more unequal alternative offer for initially advantaged (disadvantaged) party, *E_A_*(*E_B_*) is the payoff of the more equal alternative offer for the initially advantaged(disadvantaged) party, and *α* is the inequality aversion parameter that captures the decision weight of the offer inequality level. Since in the current paradigm, the two alternative offers have the same payoff sum (*I_A_* + *I_B_* = *E_A_* + *E_B_*), these two sums cancel out and the utility difference (*ΔU*) is mainly driven by the inequality difference between the two offers *ΔF* = |*I_A_* − *I_B_*| − |*E_A_* − *E_B_*|. Therefore,

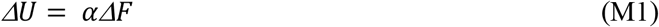

Model M2 quantified the additional effect of rank reversal aversion, on top of inequality concerns. Since our design entails that only the more equal alternative offer will reverse the relative rankings between the two parties, participants who are averse to rank reversal will thus devalue the utility of the more equal offer (*U*(*Equal*)). We thus included one discounting parameter *δ* to capture rank reversal aversion for the more equal offer, and this parameter can be seen as a measure of individuals’ bias for more equal but rank-reversing offers irrespective other motives (For detailed expositions, see Materials & Methods and Supplementary Methods sections):

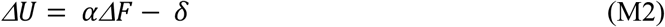

Capturing the effects of harm aversion, on top of inequality concerns and rank reversal aversion, requires somewhat more complex model assumptions, which we embedded in the following models that assumed different strategies of devaluing harms. In model family M3 (M3a - M3c), we assumed that people would devalue the utility of the alternative offer by the amount of money transferred from the initially advantaged party to the disadvantaged party. Therefore, in addition to the difference in inequality level (*ΔF*) and rank reversal, participants also considered and weighted the difference in the amount of money transferred across the two parties between the two offers (*ΔT*):

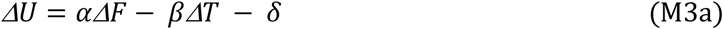

where *ΔT* is the difference in the amount of money transferred across the two parties between the two alternative offers (i.e., *ΔT* = *I_A_* − *E_A_*), and *β* is the harm aversion parameter that captures the subjective cost associated with taking money away from the advantaged.

In model family M4 (M4a - M4c), we considered that people may be not averse to transfer money away from the advantaged party as long as this transfer decreases the initial inequality level; but that they are averse to transferring more money than necessary to achieve a more equal payoff. The more unequal alternative offer decreased the inequality level but did not reverse the relative rankings or transferred more money than necessary, so that the utility of the more unequal offer was only devalued by inequality level. However, the more equal alternative offer in the critical condition transferred more money than necessary to reach the same equality level as the more equal offer itself (or the counterpart offer in the baseline condition). As the example shown in Figure 1B, Offer 2 in the baseline condition (Person A: Ұ10, Person B: Ұ8) transferred Ұ5, and the counterpart Offer 2 in the critical condition (Person A: Ұ8, Person B: Ұ10) transferred Ұ7 away from Person A’s initial endowment. Therefore, the two types of Offer 2 achieved the same equality level (i.e., absolute payoff difference between parties), but the one in the critical condition had to transfer “extra” money relative to the one in the baseline condition (i.e., Ұ7 - Ұ5 = Ұ2 in the above example). We thus considered this “extra” transferred money as unnecessary loss or harm for the initially advantaged party (i.e., H, weighted by the harm aversion parameter β). Note that, with this assumption, it is not necessary for participants to explicitly memorize and compare the two counterpart equal offers between the critical and baseline condition. Instead, they only needed to compare the more equal offer with a counterfactual offer in which the payoffs were flipped between the two parties. Therefore, in our paradigm, this “extra” transferred money equals the payoff difference in the more equal offer (i.e., *H* = *E_B_* − *E_A_*). We referred to this amount of “extra” money (*H*) as harm signals in following analyses. Therefore, Model M4a is as follows:

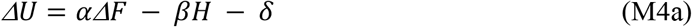

where *β* is the harm aversion parameter that captures the subjective cost of taking more money than necessary from the advantaged party.

For the model families M3 and M4, models within the same family shared the same way of calculation of harm but assumed different types of devaluations of harm and rank reversal. Specifically, M3b and M4b did not consider rank reversal, and M3c and M4c assumed that the harm aversion parameter weighed effects of both the magnitude of harm and rank reversal. For detailed expositions of the above models and other alternative models, please see Materials & Methods and Supplementary Methods sections.

To evaluate which combination of motives was most parsimonious to capture behavior, we performed formal comparisons of all the models using the Bayesian information criterion (BIC). This revealed that model M4a outperformed all other models (Table S7), suggesting that people indeed consider all three motives (inequality aversion, harm aversion, and rank reversal aversion) during wealth redistribution. Moreover, the specific form of this model entails that people only consider offers harmful if these entail taking more money than would be necessary to reach a given equality level without reversing the initial rankings (M4a). To ensure that the winning model indeed can separate identify inequality aversion (*α*), harm aversion (*β*), and rank reversal aversion (*δ*) in the critical condition, we performed parameter recovery analyses. The results confirmed that the three parameters could be recovered reliably and independently of each other (Figure 2A), indicating that our paradigm and model could clearly uncouple the effects of these different motives on individuals’ redistribution behaviors. Simulation analysis of the winning model (M4a) also showed that the probability of more equal choice varied with all the three parameters (i.e., *α, β*, and *δ*, Figure S3), indicating that different motives independently affect individuals’ redistribution decisions.

**Figure 2.**
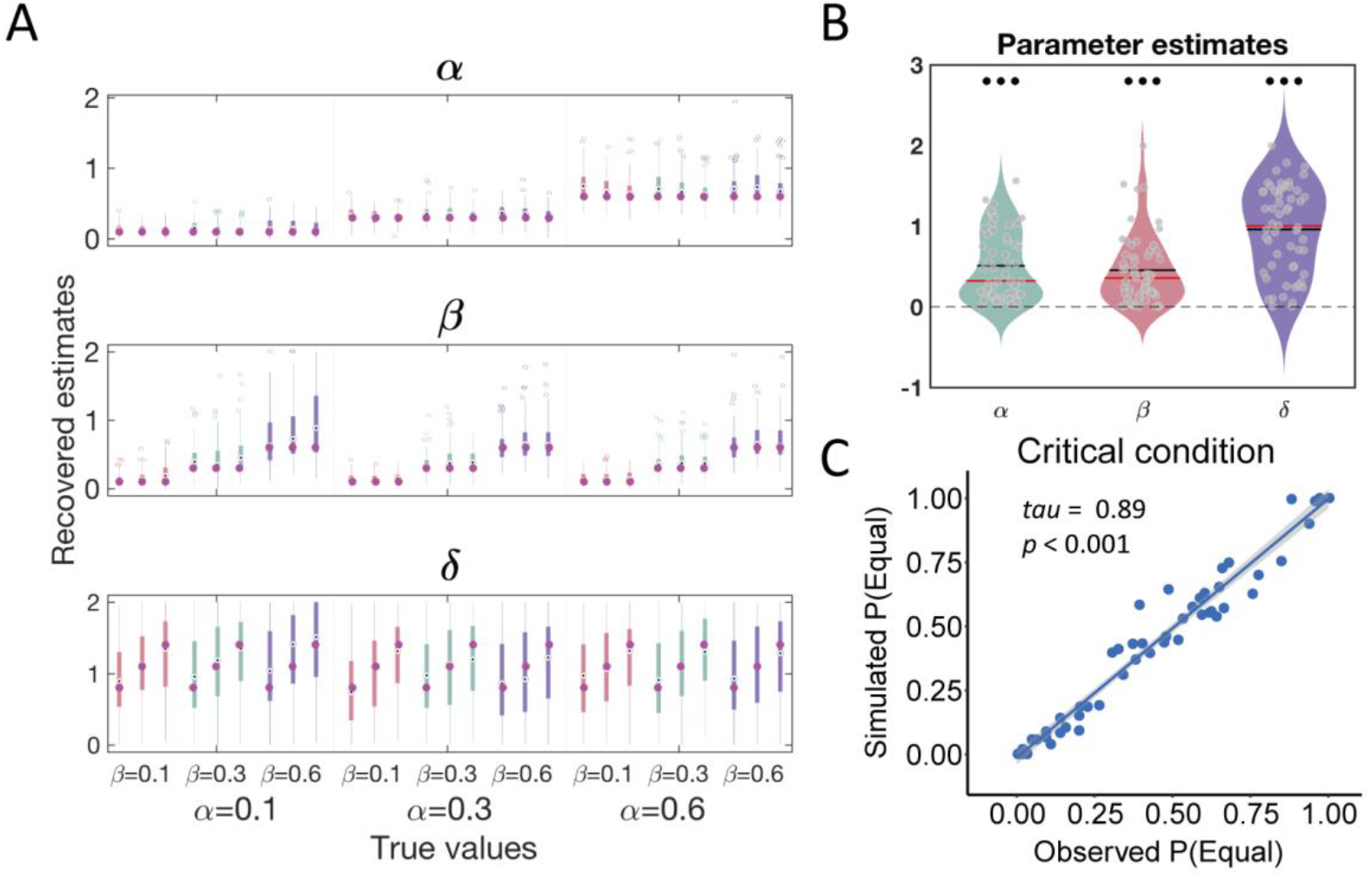
Computational modelling results. **(A)** Parameter recovery results. Box plots show that the three parameters can be recovered reliably and independently of each other, indicating that our paradigm and model can clearly uncouple the effects of different motives on individuals’ redistribution behaviors. To do this analysis, we generated 27 datasets using all combinations of three plausible values for each parameter (α: 0.1, 0.3, 0.6; β: 0.1, 0.3, 0.6; δ: 0.8, 1.1, 1.4). The boxes represent the distributions of the recovered parameters from 150 simulation sets of each combination of parameters. Each column corresponds to one type combination. Purple dots show the true values of the parameters. The recovered parameter distributions are around the true parameter values, and only vary with the true value of the parameter itself, and not with the other parameters. **(B)** Parameter estimates of the winning model M4a show that individuals exhibit inequality aversion (α), harm aversion (β), and rank reversal aversion (δ) during wealth redistribution. Violin plots show the distribution of the parameters. Black lines indicate the means, and red lines indicate the medians of the parameters at group level. Each grey dot represents one participant. **(C)** Model simulation results. The scatter plot shows a strong correlation between observed probability to choose more equal offer and model simulated probability to choose more equal offer based on model M4a in the critical condition, which suggests a good recovery performance of model M4a.

In line with model-free analyses, analyses of the winning model’s parameters confirmed again that participants’ redistribution behaviors in the critical condition were driven by inequality aversion, harm aversion, and rank reversal aversion: Specifically, participants weighed the inequality difference between the two alternative offers positively (*α* = 0.51 ±0.06, *t*(56) = 8.90, *p* < 0.001, Cohen’s d = 1.18), devalued the more equal offer by the extra harm for the initially advantaged party (*β* = 0.45 ±0.06, *t*(56) = 7.83, *p* < 0.001, Cohen’s d = 1.04), and valued rank reversal negatively (*δ* = 0.96 ±0.07, *t*(56) = 13.23, *p* < 0.001, Cohen’s d = 1.75, Figure 2B). In line with our expectations, greater inequality aversion (*α*) was associated with higher probability to choose the more equal offer (*tau* = 0.74, *p* < 0.001, Figure S4 left panel). By contrast, greater harm aversion (*β, tau* = - 0.27, *p* = 0.004) and greater rank reversal aversion (*δ, tau* = - 0.63, *p* < 0.001) were associated with higher probability to choose the more unequal offer (Figure S4 middle and right panels). Moreover, model simulation analyses showed that the choice probabilities predicted by the winning model indeed captured the observed choice probabilities well (*tau* = 0.89, *p* < .001, Figure 2C), and cross-validation prediction analyses revealed that the winning model had the highest prediction accuracy (Table S7). All these results confirmed that the winning model performed well at capturing the three latent motives underlying the observed decisions. Interestingly, correlation analyses revealed that inequality aversion (*α*) and rank reversal aversion (*δ*) were negatively correlated with each other (*tau* = - 0.62, *p* < .001, Figure S5 right panel). Given the posterior predictive checks and parameter recovery results, this correlation is very unlikely due to poor model performance and much more likely to indicate a true negative relation in these two motives.

### Neuroimaging results

As behavioral and modelling analyses have suggested that participants jointly consider inequality aversion, harm aversion, and rank reversal aversion to make redistribution decisions, we investigated how each of these motives is implemented in the brain with neuroimaging data. First, we clarified how these motives (e.g., equality and harm signals) are represented in the brain. We defined equality and harm signals based on the winning model (M4a). Next, we examined how these motives guide behavior, by investigating how the corresponding activity is modulated and interacts in line with how much the motive is evident in the behavioral effects. That is, we tested whether neural responses to equality signals differed across conditions and were modulated by activity in regions related to harm processing or rank reversal, in a manner that correlates with the observed behavioral effects.

#### Striatum represents equality and drives more equal choice

We first examined how signals associated with inequality aversion and harm aversion were represented in the brain, by constructing a general linear model 1 (GLM 1) containing parametric regressors corresponding to equality in both conditions and harm (*H*) in the critical condition while controlling statistically for the inequality in the initial offer (see Materials & Methods). In our parametric fMRI analyses, we defined equality signals as −*ΔF* = |*E_A_* − *E_B_*| − |*I_A_* − *I_B_*| so that higher equality values corresponded to smaller differences in inequality between the two alternative offers. The rationale for this definition was that people may focus more on equality signals and use this evidence to guide their decisions when the equality differences between offers is smaller. By contrast, when other motives conflict with equity-pursuing motives, responses to smaller equality differences may be inhibited, and motives to avoid harm may take over to guide decisions. We did not differentiate neural signals of rank reversal in this GLM, since the effect of rank reversal derived from contrasting between the critical and baseline condition may be confounded by other effects, such as the amount of transferred money and response bias. We focused this analysis initially on regions-of-interest (ROIs) in the striatum and ventromedial prefrontal cortex (vmPFC), which are known to be critically involved in equality processing (Tricomi et al., 2010).

Our analyses confirmed that activity in striatum, but not vmPFC, was related to equality. Specifically, activity in bilateral caudate/putamen (left MNI peak coordinates: [−18, 11, 1], k =111, max t-value = 3.64; right MNI peak coordinates: [15, 20, −5], k = 76, t = 3.55) varied parametrically with equality (− Δ *F*) in the baseline condition (Table S8, Figure 3A), but not in the critical condition. A direct comparison between striatal activity across both conditions further confirmed a more positive parametric effect of equality in the baseline than critical condition (MNI peak coordinates: [6, 14, −5], k = 45, max t-value = 4.01, Figure 3B and Figure S6 for a visualization of this effect), and this effect was also confirmed in a whole brain analysis (Table S8). The absence of striatum responses to equality in the critical condition may be due to competition between inequality aversion and the other motives that are stronger in this condition, a possibility that we further tested in analyses described later.

**Figure 3.**
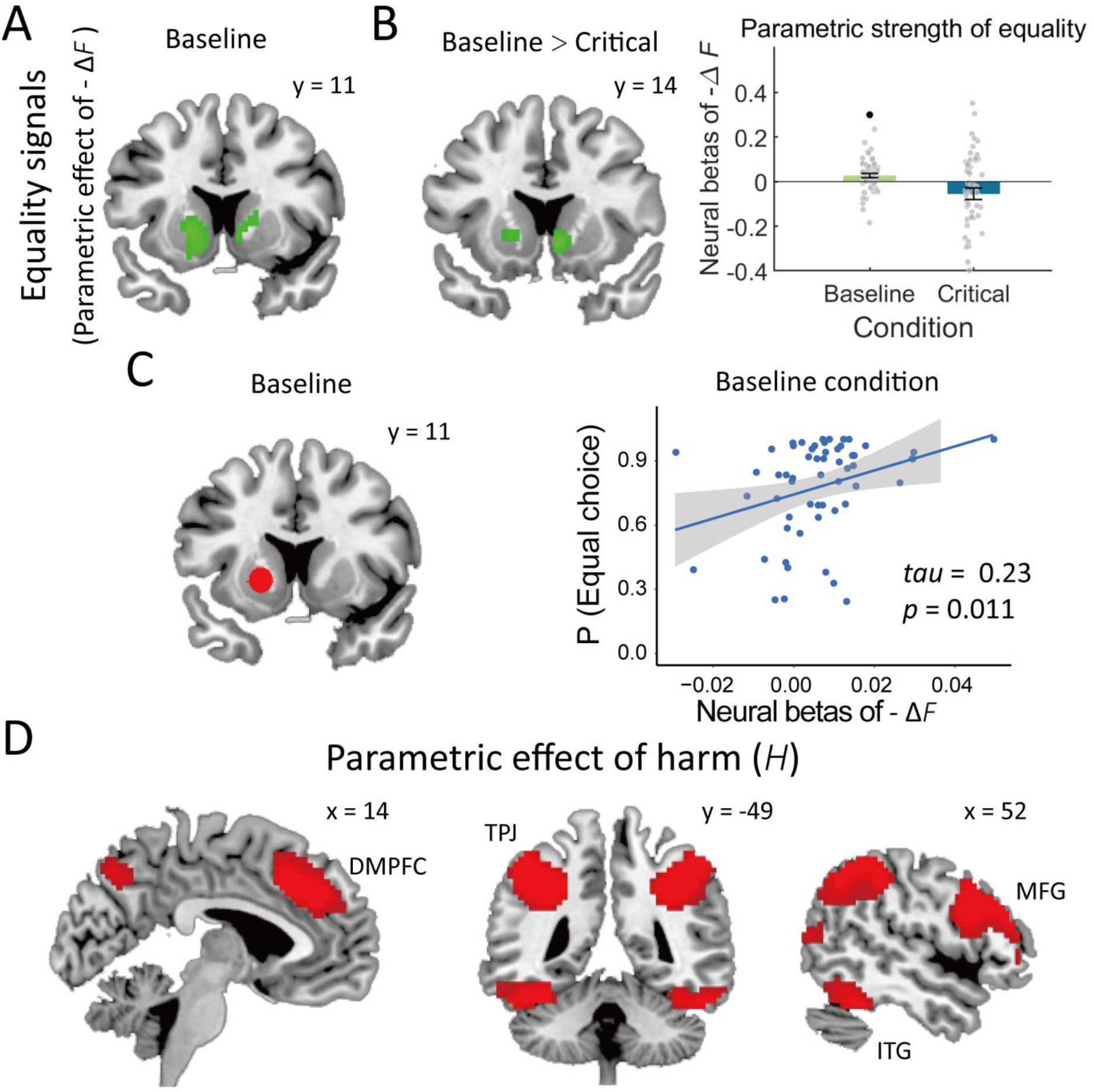
Neural representations of equality and harm signals. **(A)** Activity in caudate/putamen was associated with equality signals (-ΔF) in the baseline condition. **(B)** More positive parametric strength of equality signal in striatum in the baseline condition than in the critical condition (Left panel). For purpose of visualization, neural estimates of the significant cluster were extracted from both conditions (Right panel). Each dot represents one participant, and error bars indicate the SEMs. •, *p* < 0.05. **(C)** Scatter plot shows a correlation between the parametric strength of equality signal in the striatum (a region with the center of the peak MNI coordinates: [−12, 10, −6] in the “striatum” mask from Neurosynth) and individuals’ probability to choose the more equal offer in the baseline condition, suggesting that people whose striatum is more sensitive to equality have a stronger preference for more equal distribution. **(D)** Parametric effects of harm to the advantaged party in the critical condition. Activity in DMPFC, TPJ, MFG, and ITG increased with the extent of harm to the advantaged party, suggesting stronger processing of harm signals in these brain regions. Significant clusters are thresholded at voxel-wise *p* < 0.001 uncorrected and cluster-wise FWE corrected *p* < 0.05. Correlation result in (C) is thresholded at *p* < 0.05 FWE, small volume correction with voxel-level *p* < 0.005.

Our analyses showed that vmPFC was not involved in equality processing. However, consistent with prior studies (Bartra et al., 2013; Clithero and Rangel, 2014), GLM 2 showed that this area (MNI peak coordinates: [3, 56, −14], max t-value = 2.76, cluster-wise *p* (FWE-SVC) = 0.045, with ROI in Bartra et al., 2013) was instead involved in representing the model-predicted value of the chosen option. This finding provides neural validation of our computational behavioral model. We also constructed GLM 1a in which only the equality signals (i.e., − Δ *F*) were included as parametric regressors in both conditions. This confirmed the same results as GLM 1 (Table S9, Figure S7), but showed that harm signals (i.e., *H*) in GLM 1 did not compete for variance against equality difference (i.e., − Δ *F*).

Given that striatum was involved in signaling equality in the baseline condition, we examined whether activity in this area can bias behavior in line with inequality aversion. A correlation analysis across individuals showed that greater sensitivity to equality signals (i.e., more positive parametric estimates of − Δ *F*) in putamen (MNI peak coordinates: [−18, 11, −2], k = 6, max t-value = 2.79, cluster-wise *p* (FWE) = 0.037, small volume correction (SVC)) was indeed associated with a significantly higher probability to choose the more equal offer in the baseline condition (Figure 3C) but not in the critical condition (Figure S8). Whole brain analyses revealed that no other region correlated with individuals’ choice in either baseline or critical condition. Taken together, these findings implicate that, in situations where inequality aversion is the main motive guiding behavior, the striatum plays a critical role in processing equality and biasing redistribution behaviors in line with these concerns.

#### Cortical regions involved in signaling harm

In the critical condition, no region was associated with equality processing, but whole-brain analyses of GLM 1 showed that activity in several brain areas correlated with the harm signals related to the more equal option. These areas comprised dorsomedial prefrontal cortex/anterior cingulate cortex (DMPFC/ACC), inferior frontal gyrus (IFG), middle frontal gyrus (MFG), bilateral temporoparietal junction (TPJ), inferior temporal gyrus (ITG), and fusiform gyrus (Figure 3D, Table S8). Thus, these areas could either represent the strength of the harm aversion motive by itself, or they could be involved in processing/resolving the conflict between concerns about inequality and harm. The latter interpretation may be in line with previous findings that DMPFC/ACC, IFG, and MFG are often activated during cognitive control, conflict resolution, or behavioral adaptation (De Wit et al., 2006; Oehrn et al., 2014; de Kloet et al., 2021); and that TPJ is thought to be involved in mentalizing and perspective taking (Badre and Wagner, 2004; Van Overwalle, 2009; Oehrn et al., 2014; Hill et al., 2017). However, none of the neural effects in these areas were associated with the strength of behavioral harm aversion or inequality aversion, or the probability to choose the more equal offer in the critical condition. This motivated us to further examine whether and how the strength of the different motives was instead represented by interactions between the different neural systems involved in representing harm and equality.

#### DMPFC, as a region signaling harm, decreases neural sensitivity to equality in striatum

We had observed weaker inequality aversion, and dampened striatal sensitivity to equality, in the critical condition. These findings suggest that behaviorally relevant neural equality signals may not be represented invariably across different contexts, but may in fact be suppressed in situations where they conflict with harm signals. If this “conflict inhibition” scenario held true, then we should be able to observe that the reduction in striatal equality in the critical condition should relate to the strength of neural representations in harm-processing regions.

To test this hypothesis, we constructed a measure of neural sensitivity to equality in GLM 3 (for details, see Materials & Methods section) and identified regions that showed functional connectivity with the striatum in a way that related to the behavioral expression of harm-related inhibition of equality concerns. To do so, we subtracted the estimates of striatal activity during trials with high versus low equality signals (i.e., Beta (high -ΔF) - Beta (low -ΔF)) (Figure 4A) as our measure of neural equality signals (Figure 4B) and performed PPI analyses with the striatum as the seed region (i.e., a 6-mm radius sphere region centered at the peak coordinates of [−12, 10, −6]). As psychological context, we also selected the equality contrast (i.e., high -ΔF > low -ΔF) to examine how striatal connectivity covaries with equality in both conditions. This revealed that dorsomedial prefrontal cortex (DMPFC, MNI peak coordinates: [0, 44, 40], k = 579, max t-value = 5.09) was functionally connected with striatum more strongly for high equality signals (high - Δ F) in the critical condition (Figure 4C left panel). Consistently, PPI analyses based on GLM 1 (parametric analyses) confirmed that the same DMPFC–striatum connectivity was stronger for higher equality signals (high - Δ F) (MNI peak coordinates: [−3, 44, 43], k = 70, max t-value = 3.22, voxel-wise *p* < 0.005, cluster-wise *p* (FWE-SVC) = 0.006). Importantly, the DMPFC region identified here largely overlapped with the DMPFC region involved in signaling harm to others identified in GLM 1 (Figure 4C left panel). A whole-brain comparison also confirmed this equality-related effect on the DMPFC-striatum connectivity in the critical than baseline condition (MNI peak coordinates: [3, 50, 34], k = 40, max t-value = 3.82, cluster-wise *p* (FWE-SVC) = 0.003, Figure 4C right panel).

**Figure 4.**
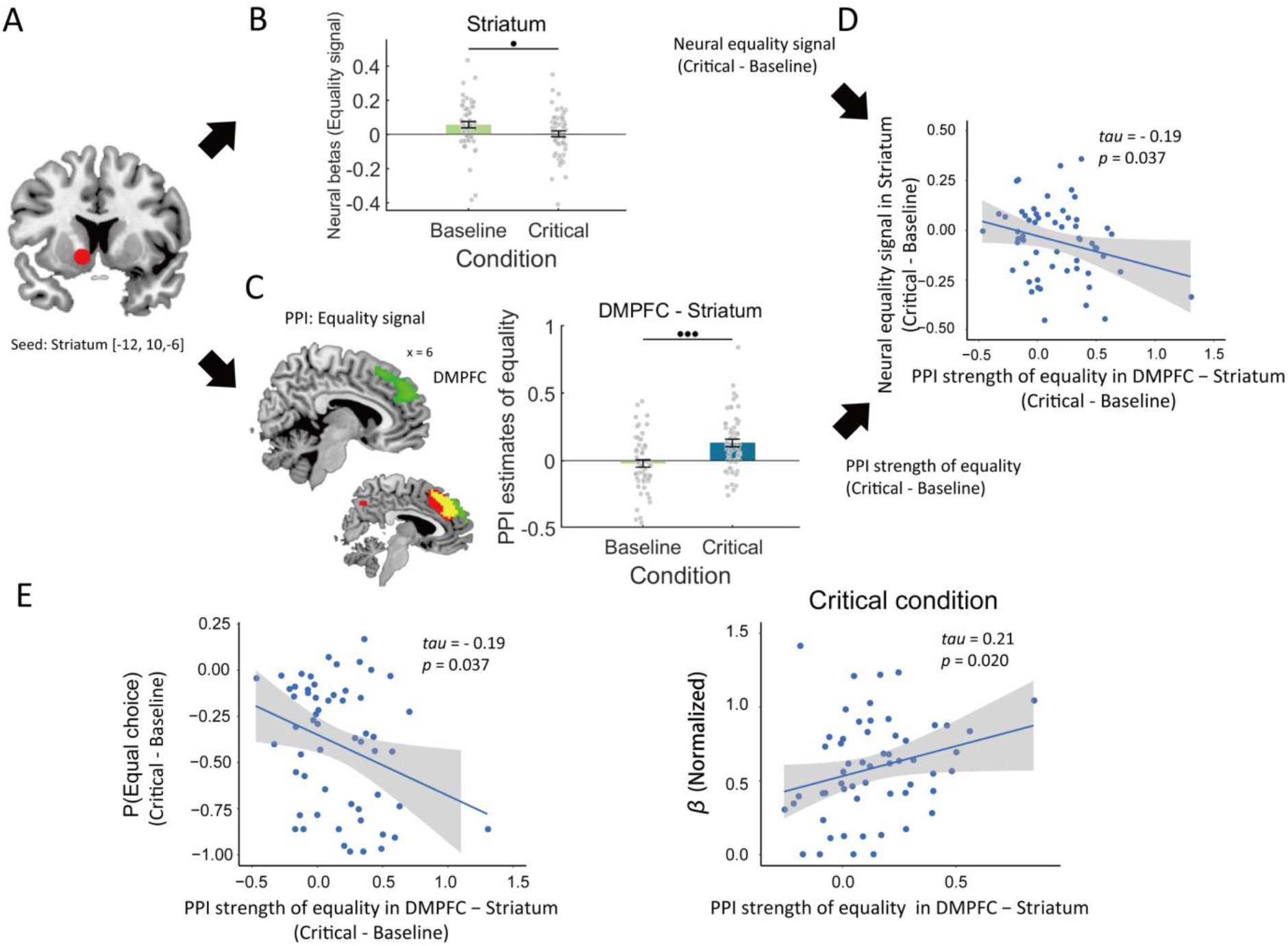
Stronger DMPFC-striatum connectivity associated with weaker neural equality signals in striatum, and behavioral effects. **(A)** We focused the context-dependent analyses on striatum region, a region with the center of peak MNI coordinates: [−12, 10, −6] in the “striatum” mask from Neurosynth. **(B)** We defined the neural equality signal as the difference in the neural estimates of striatum between high -ΔF (i.e., -ΔF = −2 and −4) and low Δ F (i.e., -ΔF = −6 and −8). Neural equality signals within the ROI showed stronger equality sensitivity in the baseline than critical condition. **(C)** PPI analysis was performed with the same striatum region as the seed to examine the connectivity strength in the contrast of “high -ΔF > low -ΔF”. It suggested a stronger DMPFC-striatum connectivity effect of equality specifically in the critical condition (left panel, DMPFC region in green), and this DMPFC region largely overlapped with the DMPFC region identified to process harm signals in GLM 1 (in red). The yellow area is the overlapping region. Post-hoc analyses further confirmed a stronger effect of equality on PPI strength in the critical condition than in the baseline condition. For visualization, we extracted the contrast value of the PPI regressors of the baseline and critical conditions within the significant cluster (right panel). **(D)** Scatter plot shows that a stronger DMPFC-striatum PPI strength of equality is associated with a lower striatum neural sensitivity to equality in the critical condition compared with the baseline condition, indicating that the lower neural responses to equality in the critical condition may be due to a stronger inhibition from DMPFC, which encodes and conveys the harm signals. **(E)** Scatter plots show that stronger equality-related DMPFC-striatum PPI connectivity is associated with a lower probability of choosing the more equal option (left panel), and with greater harm aversion (β) (right panel), in the critical relative to baseline condition. Each grey dot in (B) and (C) represents one participant, and error bars represent SEMs. •••, p < 0.001; ••, p < 0.01; •, p < 0.05. Significant clusters are thresholded at voxel-wise *p* < 0.001 uncorrected and cluster-wise FWE corrected *p* < 0.05.

Next, we tested for the critical condition whether the weaker striatum sensitivity to equality and the dampened tendency for equal choice was related to a stronger effect of equality signals on DMPFC-striatum connectivity. To do this, we correlated the differences between the critical and baseline condition in the strength of equality-related DMPFC-striatum connectivity with the differences in neural equality sensitivity in striatum. The analyses revealed a significantly negative correlation between the two variables (*Kendall’s tau* = - 0.19, *p* = 0.037, Figure 4D), showing that stronger equality-related DMPFC-striatum connectivity was indeed associated with lower neural equality sensitivity in the striatum. Moreover, greater equality-related DMPFC-striatum connectivity estimates were also associated with a lower probability of more equal choice in the critical relative to the baseline condition (*tau* = - 0.19, *p* = 0.037, Figure 4E left panel), and with stronger harm aversion (*β, tau* = 0.21, *p* = 0.020, Figure 4E right panel). These findings show that stronger DMPFC-striatum connectivity relates to greater harm aversion, lower neural sensitivity to equality in the striatum, and less equal distribution choices in situations where equity-seeking motives (i.e. inequality aversion) are in conflict with harm avoidance motives (i.e. harm aversion and rank reversal aversion).

#### Activity in cognitive control and harm-related areas during unequal choices

We also more directly examined how specific choice outcomes related to the trial-by-trial strength of neural motive representation, in terms of both regional activity and connectivity. This appears relevant for understanding what neural processes may lead individuals who are averse to inequality to nevertheless choose the more unequal offer on a specific trial. One possibility in that regard is that neural representations related to harm aversion or rank reversal aversion may be stronger for such choices. To address this issue, we constructed GLM 4 that modelled neural responses with respect to specific choices in each condition (i.e., four onset regressors in total: equal choice or unequal choice in the baseline condition and critical condition) and performed PPI analyses based on this GLM.

Our analyses revealed that in the baseline condition, activity in MFG, IFG, and TPJ was increased when participants chose the more unequal offer (contrast: unequal choice > equal choice). No region was activated for the reverse contrast (contrast: equal choice > unequal choice) in the baseline condition, or in any of the two contrasts in the critical condition (Table S10). Statistical interaction analyses between condition and choice (i.e., contrast: Baseline_(unequal choice – equal choice)_ > Critical_(unequal choice – equal choice)_) confirmed that activity in right MFG (MNI peak coordinates: [36, 32, 34], k = 41, max t-value = 3.44, cluster-wise *p* (FWE-SVC) = 0.004) and bilateral TPJ (right MNI peak coordinates: [48, −46, 34], k = 49, max t-value = 3.78, cluster-wise *p* (FWE-SVC) = 0.003; left MNI peak coordinates: [−57, −46, 28], k = 2, max t-value = 3.37, cluster-wise *p* (FWE-SVC) = 0.017) were enhanced during unequal choices specifically in the baseline condition, but not in the critical condition (Figure S9). This suggests that going against equity concerns during unequal choices may involve high-level control and/or mentalizing processes implemented by MFG and TPJ.

An involvement of prefrontal and temporoparietal control processes in unequal choices was also suggested by correlation analyses of the participants’ inequality and harm aversion parameters with brain activity related to unequal versus equal choice. This showed that the strength of activity in DMPFC (Figure 5B left panel) and TPJ (Figure 5B middle panel) during unequal choices was positively associated with inequality aversion (i.e., *α*, Table S10); and that the strength of activity in putamen was positively associated with harm aversion (i.e., *β*, Figure 5B right panel, Table S10). Note that these results were robust to the exclusion of outliers (*tau(DMPFC - α)* = 0.27, *p* = 0.007; *tau(TPJ - α)* = 0.36, *p* < 0.001; *tau(Putamen - β)* = 0.29, *p* = 0.003) and to statistical control for the effect of the other two parameters (see Supplementary Results for details). Activity in these three regions did not differ between unequal and equal choice in the critical condition at group level (Figure S10). As shown in the previous GLM analyses, both DMPFC and TPJ are involved in harm signaling, and the same DMPFC region is further involved in suppressing equality signaling in the critical condition. In line with these observations, we found that activity in DMPFC and TPJ was enhanced more strongly when more inequality-averse individuals chose the more unequal offer, again implying that activity related to harm aversion as represented in DMPFC and TPJ may deter more equal distributions, in particular for people who are averse to inequality.

**Figure 5.**
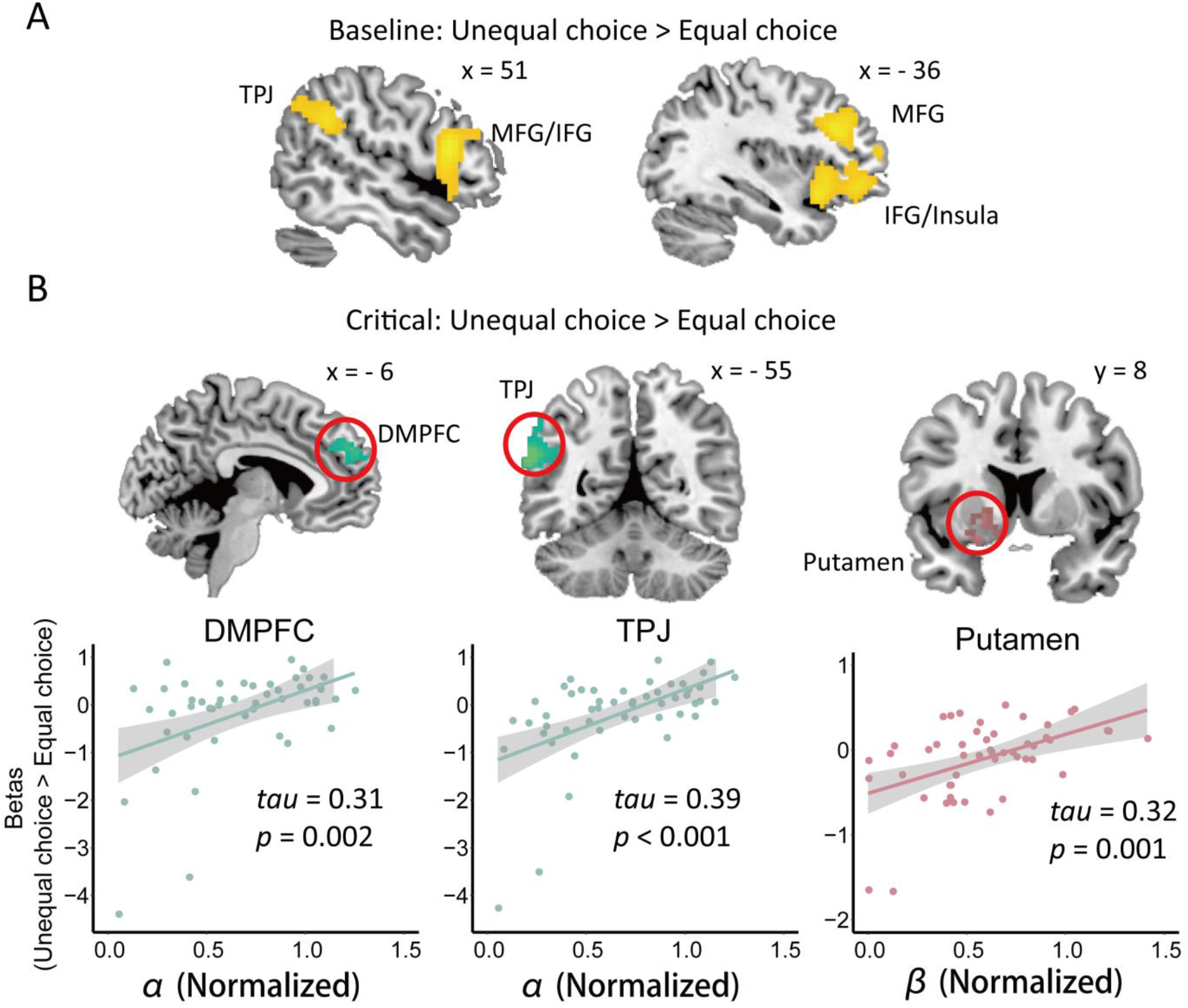
Neural responses associated with more unequal choice and linking latent motives to behaviors. **(A)** In the baseline condition, activity in MFG, IFG/Insula, and TPJ was enhanced when individuals chose the more unequal offer. **(B)** In the critical condition, activity in DMPFC (left panel) and TPJ (middle panel) was enhanced when more inequality-aversive individuals (i.e., higher α) chose the more unequal offer, whereas activity in putamen was enhanced when more harm-aversive (i.e., higher β) individuals chose the more unequal offer. For the purpose of visualization, neural estimates of the significant cluster were extracted and scatter plots show the correlation patterns (Bottom panel). Significant clusters are thresholded at voxel-wise *p* < 0.001 uncorrected and cluster-wise FWE corrected *p* < 0.05.

#### Different motives affect choice via differential patterns of network interactions

The patterns of results until now suggest that inequality and harm aversion are implemented by different neural systems, which functionally interact and inhibit one another during redistribution choice. To test more directly for the relation between choice outcome and such network interactions, we performed PPI analyses focusing on the contrast between unequal choice and equal choice in the critical condition, and considered striatum (involved in equality processing, GLM 1) as the seed region. In particular, we examined how such network interactions may be expressed in individuals with strong behavioral expression of the different motives.

We examined two possibilities in this respect. First, for individuals with stronger inequality aversion to take unequal choices, harm- or rank-reversal-related neural activity may need to be strongly recruited to inhibit striatum responses to equality signals. Thus, in inequality-averse individuals, we should see stronger activity in harm- or rank-reversal-related neural systems and stronger inhibitory connectivity with striatum during more unequal choices (see also Crockett et al., 2017; Loewke et al., 2021 for similar suggestions). Alternatively, individuals with strong harm- and rank-reversal aversion may exhibit more intense processing of the corresponding information and thus enhanced communication between the regions involved in these motives, reflecting more neural evidence about potential harm and other consequences such as rank reversal during more unequal choices.

To examine these possible neural underpinnings of unequal choices, we took the striatum region (a sphere with 6-mm radius centered on peak MNI coordinates of [−18, 11, −2]) involved in equality processing and equal choice in the baseline condition as the seed region in the PPI analyses. By doing this, we can identify whether other systems interact specifically with (e.g., inhibit) the striatum during unequal choices. This analysis confirmed that the connectivity strength between striatum and right IFG (MNI peak coordinates: [57, 26, 13], *k* = 113, max *t-value* = 5.34, Table S12) increased more strongly in people with greater inequality aversion when they chose the more unequal versus equal offer (i.e., normalized α, *tau* = 0.40, *p* < 0.001, Figure 6A & 6B top panel). This suggests that the striatum receives stronger inhibition from the IFG when more inequality-aversive individuals choose the more unequal offer. Moreover, the connectivity strength between striatum and superior frontal gyrus (SFG, MNI peak coordinates: [−21, −1, 49], *k* = 165, max *t-value* = 5.46, Table S12) increased more strongly in people with greater rank-reversal aversion when they chose the more unequal offer (i.e., δ, *tau* = 0.38, *p* < 0.001, Figure 6A & 6B bottom panel), suggesting that rank reversal aversion may inhibit equality processing via stronger inhibition signals from this area to striatum and bias the choice. It is noteworthy that although inequality aversion (i.e., α) and rank reversal aversion (i.e., δ) are negatively correlated with each other, the findings that these two motives are related to differential connectivity patterns with striatum provide evidence that they function as two different motives that independently modulate neural circuitry underlying redistribution behaviors. The correlation patterns of the above networks also held after controlling for the effect of the other two model parameters (see Supplementary Results for details).

**Figure 6.**
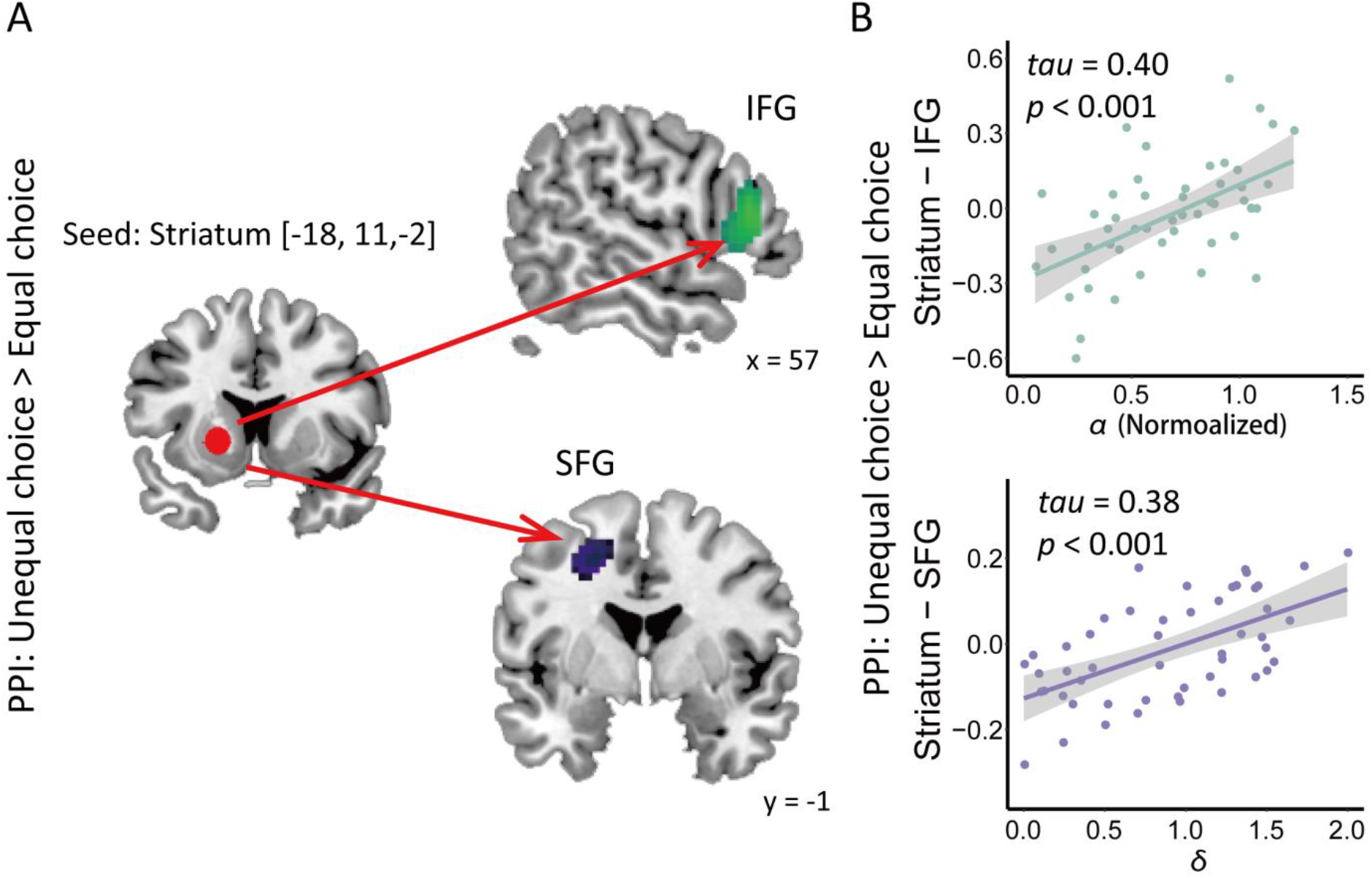
Neural networks linking different motives to redistribution decisions. (A) In the critical condition, connectivity strength between striatum and IFG was enhanced when more inequality-aversive (higher α) individuals choosing more unequal offer than more equal offer top panel), and connectivity strength between striatum and SFG was enhanced when more rank reversal-aversive (i.e., higher δ) individuals choosing more unequal offer than more equal offer (bottom panel). Neural estimates of the significant cluster were extracted and scatter plots show the correlation patterns (B). Significant clusters ware thresholded at voxel-wise *p* < 0.001 uncorrected and cluster-wise FWE corrected *p* < 0.05.

We did not observe striatal connectivity specifically associated with harm aversion in this analysis, but together with the observations of brain activity and connectivity associated with harm aversion shown in previous analyses, our findings emphasize that distinct neural pathways link different motives (inequality aversion, harm aversion, and rank reversal aversion) to redistribution behaviors, with striatal responses to equality being dampened by prefrontal areas in people with stronger aversion to inequality, harm, and rank-reversal.

## Discussion

It is widely acknowledged that increased inequality is associated with more risk-seeking behaviors, higher crime rate, and greater health problems (Choe, 2008; Pickett and Wilkinson, 2015; Payne et al., 2017). Therefore, the question of how to achieve distributive justice has become an intensively studied issue among researchers in a variety of fields, including economics, politics, philosophy, and psychology. Although influential deontological theories claim that fairness norms take precedence over other concerns (e.g., efficiency) underlying distributive justice (Rawls, 1999), empirical evidence challenges this view and suggests that other motives, such as concerns for endowment and efficiency, can undermine fairness norms and deter equal distribution (Hsu et al., 2008; Leliveld et al., 2008). However, previous studies mainly focused on how self-interest motives may run counter to inequality concerns to affect wealth distribution, and most prevailing econometric models cannot explain why individuals can prefer greater inequality when different motives conflict with each other (Fehr and Schmidt, 1999; Charness and Rabin, 2002; Tricomi et al., 2010; Saez et al., 2015; Gao et al., 2018). Although previous studies have demonstrated that harm aversion and rank reversal aversion, as two kinds of prosocial motives, are indeed involved in modulating moral decisions and redistribution decisions (Crockett et al., 2014, 2017; Xie et al., 2017), it is still largely unclear how these motives interact with inequality aversion to bias individuals’ choices.

To bridge these gaps, the current study establishes a novel redistribution paradigm and an integrated computational modelling approach to examine how conflicts between different prosocial motives bias individuals’ preferences in wealth distribution. We demonstrate that harm aversion and rank reversal aversion can substantially suppress equality processing, which is considered as the core principle of distributive justice, to prevent more equal distribution. At the neural level, we suggest that the striatum serves as a hub for signaling equality and guiding decisions in line with equality concerns; and that striatal representations of equality will be suppressed by other systems when these are in conflict with harm avoidance and rank preserving motives. Importantly, interactions between different regions of frontal cortex and striatum play critical roles to modulate different motives and lead to more unequal choices.

Our study extends economic theories of social preferences by highlighting the trade-off between multiple prosocial motives in third-party wealth distribution, and by exploring the boundaries within which inequality aversion determines wealth redistribution behavior. In the literature of third-party norms, theories often argue that people are inclined to punish norm violators in order to facilitate social norms (Fehr and Fischbacher, 2004a, 2004b; Hu et al., 2015). The current paradigm excludes the possibility of intentional violation of fairness norms, since the initially unequal distributions were generated from random draws. Given that participants still exhibit strong preferences for equal distribution in such situations, we suggest that inequality aversion, rather than motives to punish norm violation, drives redistribution behaviors as a core principle in wealth redistribution. However, importantly, we observed that people weighed equality less when it conflicted with preferences for harming others (i.e., harm aversion) or preserving initial rankings (i.e., rank reversal aversion), suggesting that equality-seeking motives (i.e., inequality aversion) are coordinated with other prosocial motives in wealth redistribution. These findings further extend influential theories of fairness norms (Fehr and Schmidt, 1999; Charness and Rabin, 2002) which mainly focused on effects of inequality aversion on distribution behaviors, and emphasize the importance of also considering other motives (i.e., harm aversion and rank reversal aversion) in econometric models, especially since conflicts between these different motives are prevalent in real life distribution decisions (e.g., taxation policy).

Harm aversion, as a critical type of moral virtue, drives people to achieve a more equal distribution by transferring as little money as possible between two parties. In line with our observation, when making moral decisions, people typically conform to the ‘do-no-harm’ principle and prefer not to benefit one party by harming another party (Baron, 1993, 1995; Crockett et al., 2014). One potential explanation for harm aversion stems from studies of morality suggesting that people are not willing to take responsibility for others’ bad outcomes when making moral decisions (Ritov and Baron, 1990; Crockett et al., 2014), as such moral responsibility will induce individuals’ anticipatory guilt emotion which proscribes people from harming others (Chang et al., 2011; Yu et al., 2014a). Therefore, in the current paradigm, taking more money away from others brings not only greater cost for the initially advantaged party but also greater cost of moral responsibility (i.e., harm aversion) for participants which will in turn dampen their motives to seek equality.

Moreover, we suggest that rank reversal aversion is another prosocial motive that discounts the utility of equality during wealth redistribution. A stable hierarchy can provide fitness advantage by facilitating social information processing (Zitek and Tiedens, 2012), satisfying individuals’ psychological need for order (Friesen et al., 2014), and enhancing intragroup cooperation and productivity (Halevy et al., 2012; Ronay et al., 2012). Therefore, it is not surprising that people prefer to preserve rather than reverse pre-existing hierarchy (Fernandez and Rodrik, 2004; Xie et al., 2017). In line with these findings, our results suggest that the reversal of initial rankings also contributes to the disutility of equality when rank preserving and equality seeking are in conflict. Together, taking advantage of computational modelling approaches, we demonstrate that in contrast to inequality aversion, harm aversion and rank reversal aversion function as two different third-party prosocial preferences to deter more equal wealth redistribution.

Our neural imaging findings first clarified how equality-related information is represented. GLM results support the hypothesis that individuals are sensitive to equality signals in the absence of any conflict but will be less sensitive to equality and base their decisions more heavily on other motives when they conflict with inequality aversion. Although previous studies have proposed that the striatum signals rewarding aspects of equality-related distributions (Hsu et al., 2008; Tricomi et al., 2010; Hu et al., 2015), it is still largely unclear which specific aspects of the distributions behavior engage the striatum and trigger the corresponding behavior – does it signal equality or other potentially rewarding aspects, such as efficiency or the other’s outcomes? While stronger activity in putamen has been previously found to be related to higher efficiency (i.e., greater overall profits) (Hsu et al., 2008), efficiency cannot account for the pattern of results in the current study since neither of the two alternative offers changed the overall profits of the distributions. An alternative plausible explanation is that striatum activity reflects dopaminergic responses in reward computation of social welfare, as it has been widely observed that stronger striatum activity is associated with charitable giving (Moll et al., 2006; Harbaugh et al., 2007), altruistic punishment to norm violation (De Quervain et al., 2004), and more equal wealth distributions (Tricomi et al., 2010; Hu et al., 2015).

Moreover, striatum was involved in arousal representations (Knutson et al., 2000, 2001). For example, stronger striatal activation was related to greater motivation of norm compliance (Spitzer et al., 2007). In the current study, smaller equality difference between the two alternative offers may require participants to base their decisions more heavily on the evidence of equality signals, and result in stronger motivation to comply with fairness norms for them, which is manifested by enhanced striatal activity. Together with the finding that greater sensitivity to equality in putamen was related to higher probability to choose the more equal offer, our results suggest that striatum not only reflects processing of equality signals but also promotes fairness norm compliance.

Importantly, representations of equality in striatum were only observed in the baseline condition, and this striatal signaling of equality was suppressed in the context with conflicts between motives (i.e., critical condition). Moreover, stronger connectivity between striatum and DMPFC was associated with lower equality sensitivity in striatum, lower probability of equal choice, and higher strength of harm aversion in the critical condition. These findings help to clarify the neurocognitive mechanisms of the weighing processes of different motives, by providing a potential neural explanation for the weaker impact of equality on redistribution decisions in the critical condition: DMPFC may process harm information, convey harm aversion motive to striatum, inhibit striatum responses to equality signals, and dampen equal choice. Evidence from two different lines of research supports such a modulating role of DMPFC for the trade-off between motives. On the one hand, DMPFC, together with adjacent regions ACC, is critical engaged in conflict monitoring, conflict resolution, and action selection in a variety of cognitive tasks (Badre and Wagner, 2004; De Wit et al., 2006; Oehrn et al., 2014; de Kloet et al., 2021), which may support the resolution of conflict between different motives in the current paradigm (i.e., suppressing motive of equality pursuing). On the other hand, DMPFC is also thought to be part of the mentalizing system that supports vicarious experiences of others’ pain or beliefs (Van Overwalle, 2009; Garvert et al., 2015), which may support harm signals in the current paradigm. In line with our findings, inhibitory connectivity from prefrontal cortex to striatal value representations was also found to modulate individuals’ behaviors in other kinds of social and non-social decision making (van den Bos et al., 2014; Crockett et al., 2017). Future studies with brain stimulation may be needed to establish whether DMPFC influences on striatum are indeed causally involved in guiding redistribution behaviors under circumstances with conflicts between multiple motives.

Our results also provide crucial novel evidence for frontostriatal circuitry in redistribution decision making. The critical role of frontostriatal circuitry in decision making has been highlighted in both social and non-social behaviors (Spitzer et al., 2007; Rangel and Hare, 2010; van den Bos et al., 2014; Crockett et al., 2017). In general, striatum has been suggested to receive inputs of goal-related representations from lateral prefrontal cortex and output value signals to guide response selection to maximize reward (Balleine et al., 2007; Buckholtz and Marois, 2012; van den Bos et al., 2014). In line with these suggestions, lateral prefrontal cortices are usually implicated in either inhibiting intuitive motivations or modulating value representations that integrate information from different sources for moral and prosocial decision making (Feng et al., 2015; Crockett et al., 2017; Hu et al., 2021). Our findings further refine previous accounts of frontostriatal circuitry in moral decision making by clarifying that different prosocial motives modulate redistribution decisions through differential frontostriatal circuitries. Although our connectivity analyses cannot provide directional evidence, these results are in line with the view that prefrontal cortex is involved in inhibitory or executive control functions (Buckholtz and Marois, 2012; Feng et al., 2015). Striatum may receive signals from different sub-regions of prefrontal cortex that reflect different prosocial motives and bias final action selection based on integrated values (Lau and Glimcher, 2007; Grahn et al., 2008; Crockett et al., 2017).

Another critical contribution of our study is to clarify what neural processes underlie the modulations of different prosocial motives on redistribution decisions. Apart from processes involved in generally arbitrating between motives (i.e., DMPFC-striatum connectivity), it is also important to identify processes that bias behavior on a trial-by-trial level in line with different motives, and which may differ between people with different motive strengths. Activity in both DMPFC and TPJ were stronger when more inequality-aversive individuals chose the more unequal offer, and activity in putamen was stronger when more harm-aversive individuals chose the more unequal offer. One possibility suggested by the literature is that DMPFC and TPJ may support social cognitive processes such as mentalizing, perspective taking, inference and learning about others’ preferences (Decety and Lamm, 2007; Singer and Lamm, 2009; Van Overwalle, 2009; Garvert et al., 2015; Stanley, 2016; Ogawa and Kameda, 2020). Recent studies have further differentiated the roles of these two regions, by suggesting that while DMPFC is implicated in value-based action selection in a domain general manner (Rushworth et al., 2004; Rushworth and Behrens, 2008; Kovach et al., 2012; Boorman et al., 2013; O’Reilly et al., 2013), TPJ may be more specifically involved in processing of context-dependent social information (FeldmanHall et al., 2014; Santiesteban et al., 2015; Lee and McCarthy, 2016; Konovalov et al., 2021). Although our findings cannot provide a clear dissociation between DMPFC and TPJ, among all the regions involved in harm signaling, these two regions may be well-suited interfaces to link latent social motives to specific decisions. These findings also parallel the observation of stronger activity in TPJ for unequal choice vs equal choice in the baseline condition, which may implicate the role of TPJ in social cognitive processing irrespective of whether there are conflicts between different motives.

In general, our findings may have economic, political, and social implications (Irlenbusch and Villeval, 2015). Economists have introduced endowment effect for decades to explain individuals’ tendency to increase the subjective value of objects they own already (versus those they want to purchase), especially in the situation of goods trading (Carmon and Ariely, 2000). Forgoing one’s own good is seen as a kind of loss, and loss aversion or ownership will make it harder to give up the good (i.e., increased willingness to accept to give up) (Kahneman et al., 1991; Morewedge et al., 2009). Perhaps in analogy to the endowment effect (Weaver and Frederick, 2012), our study highlights that people are inclined to maintain initial relative rankings and to take less money away from others in wealth redistribution, considering the reversal of initial rankings and others’ monetary loss as a kind of third-party loss which proscribes actions to achieve higher equality (Xie et al., 2017). More generally, our findings may also explain resistance to reform policies that aim to promote social efficiency or reduce income inequality (Fernandez and Rodrik, 2004; Kuziemko et al., 2014). For instance, rich people in regions with more equal income distribution, whose advantaged ranks can be more easily reversed, are less supportive of redistribution than those in regions with more unequal income distribution (Dimick et al., 2016). Given that the effects of different motives (i.e., inequality aversion, harm aversion, and rank reversal aversion) are scientifically validated in the current study, this may help to develop better taxation policies by taking these motives into account when designing measures to reduce social inequality on the one hand and satisfy people in different income groups who pursue different motives on the other hand.

To conclude, the current study provides a neurocomputational account of the trade-off between multiple prosocial motives underlying resource distribution. Our findings suggest that in addition to inequality aversion, harm aversion and rank reversal aversion work as two separate prosocial motives to modulate individuals’ behaviors during wealth redistribution. Moreover, our study offers neural explanations for how different prosocial motives modulate redistribution behaviors, by highlighting a crucial role of striatum in equality processing and modulation of motives on ultimate decisions. Our approach improves our understanding of cognitive and neurobiological mechanisms underlying social preferences and distributive justice and may have implications for development of reform policies to promote fairness norms and social justice.

## Materials & Methods

### Participants

Sixty-three right-handed undergraduate and graduate students were recruited in the experiment. Six participants were excluded because of either making the same decision all the time or excessive head movement (> ± 3 mm in translation and/or > ± 3°in rotation). The remaining 57 participants were aged between 19 and 28 years (mean = 21.83 SD = 1.91; 31 female). No participant reported any history of psychiatric, neurological, or cognitive disorders. Informed written consent was obtained from each participant before the experiment. The study was carried out in accordance with the Declaration of Helsinski and was approved by the Ethics Committee of the Department of Psychology, Peking University.

### Design and procedures

In the present study, we developed a novel redistribution task to assess individuals’ preferences to redistribute unequal wealth allocations. In this task, participants were first presented with a monetary distribution scheme (i.e., Initial offer: Person A: Ұ15, Person B: Ұ3) between two anonymous strangers. The initial endowment of each party was allocated unequally and randomly by computer, and participants had to choose between two redistribution options (i.e., alternative offers) which transfered a certain amount of money from the one with higher initial endowment (advantaged party) to the one with lower initial endowment (disadvantaged party, Figure 1A). Participants were informed that all of the anonymous strangers in the redistribution task had made the same effort, spent equal amount of time, and made the same contribution in another experiment, and that their decisions would determine those strangers’ final payoffs. Moreover, the strangers would only know their own final payoffs but would not know their initial endowments or the payoffs of others.

In the baseline condition, both alternative offers (e.g., Offer 1: Person A: Ұ14, Person B: Ұ4; Offer 2: Person A: Ұ10, Person B: Ұ8) were more equal than the initial offer (e.g., Person A: Ұ15, Person B: Ұ3), and both alternative offers kept the same total payoffs and the same relative rankings between the two parties as the initial offer. In the critical condition, participants were presented with the same initial offer (e.g., Person A: Ұ15, Person B: Ұ3) and the same more unequal alternative offer (e.g., Offer 1: Person A: Ұ14, Person B: Ұ4) as the baseline condition, but with a different more equal alternative offer (e.g., Offer 2: Person A: Ұ8, Person B: Ұ10). This more equal alternative offer (e.g., Person A: Ұ8, Person B: Ұ10) had the same inequality level as the more equal alternative offer in the baseline condition (e.g., Offer 2 in the baseline condition: Person A: Ұ10, Person B: Ұ8), but would reverse the initially relative advantageous/disadvantageous rankings of the two parties (Figure 1B). We matched all trials in the baseline condition with the critical condition to allow for direct comparison between the two conditions. The difference in the probability of choosing the more equal alternative offer between the baseline and critical condition was then considered as a behavioral measure of the effect of harm aversion and/or rank reversal aversion on redistribution behavior. To differentiate the effect of inequality and the amount of transferred money (i.e., harm to the advantaged party), we orthogonalized the differences in inequality and the transferred money between the two alternative offers in the critical condition (Figure 1C top panel). In addition, we included two filler conditions in which one of the alternative offers was equally distributed (e.g., Person A: Ұ9, Person B: Ұ9), and the other alternative offer either kept (filler condition 1, e.g. Person A: Ұ14, Person B: Ұ4) or reversed (filler condition 2, e.g. Person A: Ұ4, Person B: Ұ14) initially relative advantageous/disadvantageous rankings (Figure 1B).

At the beginning of each trial, a fixation point was presented at the center of the screen for 1s, then the pictures of the two anonymous strangers together with their initial endowments were presented for 3 s. Next, after a blank screen jittering from 1 to 4 s, the two alternative offers were presented. Participants needed to choose one out of the two alternative offers within 6.5 s. After a blank screen jittering from 1 to 4 s, the next trial began (see Figure 1A). The participants knew that 10 trials were randomly selected by computer to determine corresponding persons’ final payoff based on their decisions. There were 66 trials in each of the baseline and critical conditions, and 15 trials in each filler condition. The 162 trials were divided into three scanning sessions lasting ~15 minutes each. After the experiment, each participant received CNY 120 (~ USD 20) for compensation.

### Model-free analysis

We first conducted model-free generalized mixed-effects analysis to test the effects of different components on individuals’ probability to choose the more equal alternative offer. In this analysis, we pooled all the trials in the baseline and critical conditions across all participants. We considered participants’ choice as the dependent variable (more equal choice = 1, more unequal choice = 0), and included 1) the absolute value of difference in the initial endowments between the two parties (Δ Initial endowment), 2) the absolute value of difference in the inequality level between the two alternative offers (Δ Inequality), 3) differences in the amount of transferred money between the more equal alternative offer and the more unequal offer (Δ Transfer), 4) condition (critical = 1, baseline = 0), and interactions between the four variables as the predictors in the model. All these predictors were standardized before being entered into the model and considered as fixed effects, and participants were considered as a random-effect intercept term. We performed this linear mixed-effects analysis using the lme4 package in R.

### Computational modelling analysis

We established four families of computational models to formally examine how inequality aversion, harm aversion, and rank reversal aversion affect individuals’ redistribution behaviors in the critical condition. Different models held different assumptions about how people discounted the utility of the more equal alternative offer [*U*(*Equal*)] in the critical condition.

Model M1 assumed that participants’ choices are only influenced by inequality aversion. We followed the classical inequality aversion model proposed by Fehr and Schmidt (1999) in which people assign values to the outcomes of all parties but devalue the inequality they experience for any kinds of distribution.

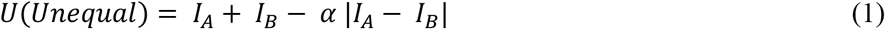

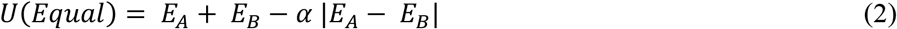

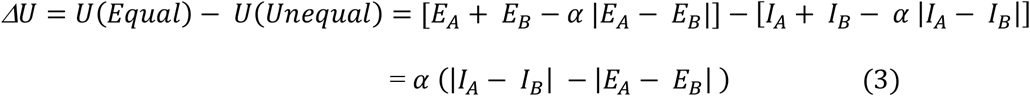

where *I_A_*(*I_B_*) is the payoff of the more unequal alternative offer for initially advantaged (disadvantaged) party, *E_A_*(*E_B_*) is the payoff of the more equal alternative offer for initially advantaged (disadvantaged) party, and *α* is the inequality aversion parameter that captures the weighing of inequality level of the offers. Since in the current paradigm, the two alternative offers have the same payoff sum (*I_A_* + *I_B_* = *E_A_* + *E_B_*), the utility difference (*ΔU*) is mainly driven by the difference in inequality level between the two offers (i.e., |*I_A_* − *I_B_*| − |*E_A_* − *E_B_*|). We refer to this inequality difference as *ΔF* (*ΔF* = |*I_A_* − *I_B_*| − |*E_A_* − *E_B_*|). Therefore,

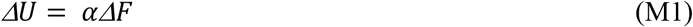

Model M2 quantified additional effects of rank reversal aversion on top of inequality concerns. We included one discounting parameter *δ* to capture rank reversal aversion for the more equal offer (For detailed expositions, see Supplementary Methods):

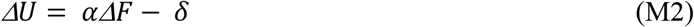

In Model M3a to M3c, we assumed that participants would first evaluate the inequality level of the two alternative offers. Then, we assumed that people would discount the utility of the alternative offer by the amount of money transferred from the initially advantaged party to the disadvantaged party. To reach the more equal offer, participants need to transfer a larger amount of money from the initially advantaged party to the disadvantaged party than to reach to the more unequal offer. Therefore, we assumed that in addition to the difference in inequality level (*Δ*F) and rank reversal, participants would also consider the difference in the amount of money transferred between the two parties (*Δ*T):

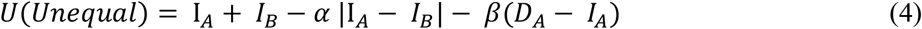

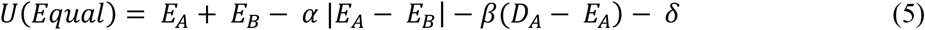

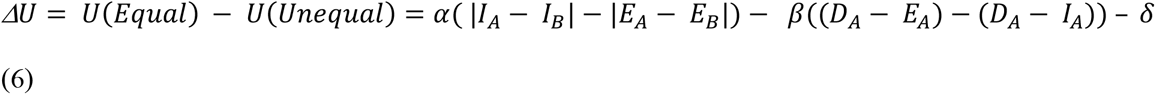

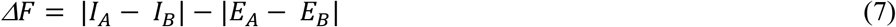

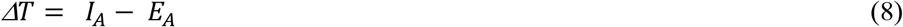

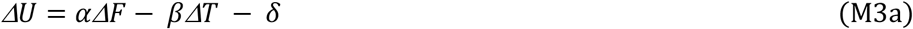

where *D_A_*(*D_B_*) is the payoff of the initial offer for advantaged (disadvantaged) party, *ΔF* is the difference in inequality level between the two alternative offers as above models, and *ΔT* is the difference in the amount of money transferred across the two parties between the two alternative offers. *α* and *δ* are still the inequality aversion parameter and rank reversal aversion parameter as control models. *β* is the harm aversion parameter that captures the subjective cost to take money away from the initially advantaged party.

In model M3b, we assumed that people did not consider rank reversal generated by the more equal offer:

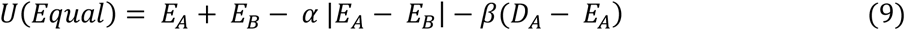

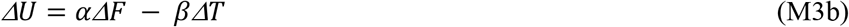

Model M3c assumed that participants discounted the utility of the more equal alternative offer for both transferred money and rank reversal, but that the harm aversion parameter captured additive effects of the transferred money and rank reversal effect:

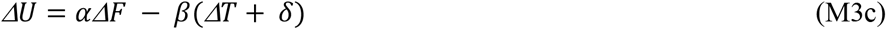

The models listed above assumed that people would discount the more equal alternative offer for the transferred money. However, it is possible that people are not averse to transfer more money as long as the transferred money can decrease the initial inequality level. Instead, they may be only averse to reach a certain equality level by transferring more money than necessary. Therefore, in models M4a to M4c, we assumed that participants would first evaluate the inequality difference between alternative offers (*ΔF*) and rank reversal. Since in the critical condition, the more unequal alternative offer decreased the inequality level of the initial distribution and did not reverse the initially relative rankings or transferred more money than necessary, the model assumed that the utility of the more unequal alternative offer was only devalued by the inequality level. For the more equal offer, we assumed that in addition to inequality level, participants would also discount *U (Equal*) for the proportion of the transferred money that exceeded the necessary amount of money that could reach the same equality level as the equal offer itself but keep the initially relative rankings (i.e., *H*), which was also considered as the amount of extra harm to the advantaged party. Therefore, model M4a is as follows:

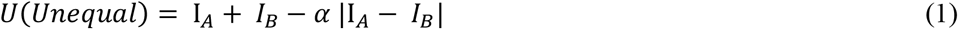

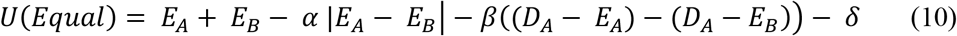

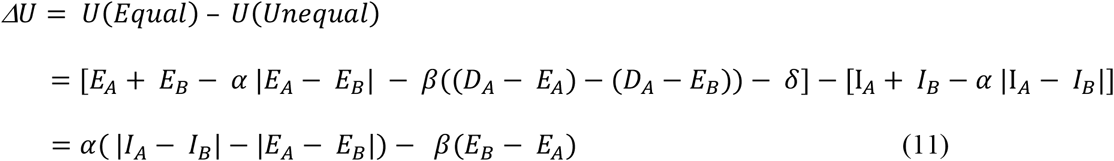

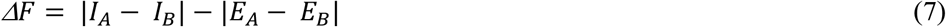

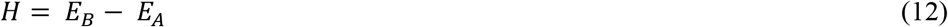

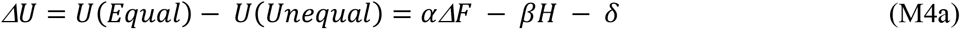

where *α* is the inequality aversion parameter that captures the weighing of inequality difference between the two alternative offers, *β* is the harm aversion parameter that captures the subjective cost of taking more money than necessary from the advantaged party (i.e., generating greater others’ loss or harm), *δ* is the rank reversal aversion parameter, and *H* is the proportion of the transferred money that exceeds the necessary amount of money that can both reach the same equality level as the equal offer itself and keep the initially relative rankings.

In models M4b and M4c, we also assumed different ways to account for individuals’ aversion to rank reversal:

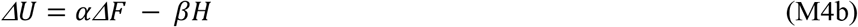

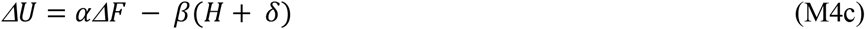

To summarize, in total, we established eight models in four families which held different assumptions about how people devalue the utility alternative offers in the critical condition. In the control models (M1 and M2), we only considered inequality aversion (M1) or additional rank reversal aversion (M2). For the two model families considering harm aversion (M3a – M3c and M4a – M4c), models within the same model set shared the same way of calculation of harm, but assumed different types of devaluations of harm and rank reversal.

Similar to the notion of the harm signals (i.e., *H*) calculated above, it is also possible that participants would devalue the more equal alternative offer only by the proportion of the transferred money that reversed the initial relative rankings (i.e., *R* = (*E_B_* − *E_A_*)/2), but not by the proportion that reduced the inequality level to the absolute equality level. This psychological component looks differently from the harm signals in M4a to M4c at the first glance but is just double the harm signal as defined above (i.e., *H* = 2 · *R*). Therefore, we did not set up a separate model for this possibility.

For all models, we calculated trial-by-trial utility differences (*ΔU*) between the two alternative offers [(*ΔU* = *U*(*Equal*) − *U*(*Unequal*)] and employed a softmax function to transform these utility differences into probabilities of choosing the more equal alternative offer:

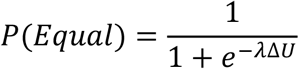

where *λ* is a free temperature parameter reflecting to what extent an individual’s decisions depend on *ΔU*.

We obtained best fitting parameters by maximizing the log likelihood of the data for each model with the MATLAB function fmincon. To avoid the optimization getting stuck in local minima, we used multiple starting points. To evaluate model fits, we calculated the Bayesian Information Criterion (BIC) (Schwarz, 1978) which rewards model parsimony to avoid overfitting:

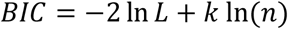

where *L* is the maximized likelihood for the model, *k* is the number of free parameters in the model, and *n* is the number of observations.

#### Parameter recovery

We performed parameter recovery to validate that the winning model can identify each parameter. We focused on how well the model can recover the three critical parameters: inequality aversion *α*, harm aversion *β*, and rank reversal aversion *δ*. Specifically, we generated 27 datasets using all combinations of three plausible values for each parameter (α: 0.1, 0.3, 0.6; *β:* 0.1, 0.3, 0.6; *δ:* 0.8, 1.1, 1.4), with the temperature parameter *λ* fixed at 1.2. For each parameter combination, we applied the set values of the three parameters to the winning model to simulate agent’s responses in the critical condition 150 times, and then re-estimated the three parameters for the simulated responses using the winning model to get 150 sets of recovered estimates. We checked how well the distributions of the recovered estimates fit with the true values of the parameters.

#### Cross-validation prediction analyses and posterior predictive checks

To further evaluate the performance of the winning model, we performed supplementary analyses. First, we did cross-validation prediction analyses by estimating model parameters with each participant’s half trials (i.e., odd-numbered trials), and simulating responses with the estimated parameters on the other half trials (i.e., even-numbered trials). Then, we calculated the cross-validated prediction accuracy by comparing simulated responses with observed responses. Second, we simulated each participant’s responses by applying their own parameters estimated from all trials to the winning model to generate 100 sets of simulated responses for each participant. Then, we calculated the probability to choose the more equal offer in these 100 sets of responses as simulated behavior and correlated it with observed probabilities of choosing the more equal offer. Note that, to avoid potential effects biased by outliers, we employed nonparametric tests (i.e., Kendall’s tau) for all correlation analyses in the current study.

### fMRI data acquisition and preprocessing

We collected T2*-weighted echo-planar images (EPI) using a GE-MR750 3.0T scanner with a standard head coil at Tongji University, China. The images were acquired in 40 axial slices parallel to the AC-PC line in an interleaved order, with an in-plane resolution of 3mm × 3mm, a slice thickness of 4 mm, an inter-slice gap of 4 mm, a repetition time (TR) of 2000 ms, an echo time (TE) of 30 ms, a flip angle of 90°, and a field of view (FOV) of 200mm × 200 mm. We used Statistical Parametric Mapping software SPM12 (Wellcome Trust Department of Cognitive Neurology, London, UK) which was run through Matlab (Mathworks) to preprocess the fMRI images. For each session, the first five volumes were discarded to allow for stabilization of magnetization. For the remaining images, we first performed slice-time correction to the middle slice, then realigned the images to account for head movement, spatially re-sampled the images to 3 × 3 × 3 isotropic voxel, normalized them to standard Montreal Neurological Institute (MNI) template space, and finally spatially smoothed the images using an 8-mm FWHM Gaussian kernel. Data were filtered using a high-pass filter with 1/128 Hz cutoff frequency.

### General linear model (GLM) analyses

We constructed the following GLMs to address specific questions. First, we built GLM 1 to examine how signals of equality and harm to others were represented in the brain during wealth redistribution in different conditions. GLM 1 included the regressor corresponding to the onset of alternative offer presentation in each condition separately (i.e., baseline, critical, and filler). The duration for these events were equal to the time form onsets of the alternative offer presentation to the time points of offsets. Moreover, GLM 1 included the trial-wise equality difference between the two alternative offers (− Δ *F*) as the parametric modulator for the alternative offer events in the baseline condition, and both the trial-wise equality difference between the two alternative offers (− Δ *F*) and trial-wise harm of the more equal alternative offer (*H*) as the parametric modulators for the alternative offer events in the critical condition. To identify neural correlates that reflected the signals of equality and harm irrespective of participants’ choices, we examined the following contrasts: ‘Baseline: equality signal (-Δ *F*)’, ‘Critical: equality signal (− Δ *F*)’, and ‘Critical: harm signal (*H*)’, respectively. To identify neural correlates that reflected the difference in the signal of equality difference between the baseline and critical condition, we examined the contrast of ‘Baseline: equality signal > Critical: equality signal’. Significant results are reported at a cluster-wise FWE corrected *p* < 0.05 (cluster-forming threshold voxel-wise *p* < 0.001 uncorrected) throughout all the analyses unless otherwise noted.

To confirm that the weaker parametric effect of equality signals in striatum for the critical condition was not because the parametric modulator of harm signals in the critical condition captured the variance accounted by equality signals, we established GLM 1a in which only trial-wise equality signal (- Δ *F*) was included as the parametric modulator for the alternative offer events in both conditions. Therefore, in the GLM 1a, the alternative offer onsets in the baseline and critical condition have the same parametric modulator (i.e., equality signal (- Δ*F)*).

To examine the neural response patterns of the parametric effects identified in GLM 1, we generated GLM 1b in which trials were divided into 4 levels of equality signals (i.e., −8, −6, −4, and −2) and 4 different alternative offer onset regressors corresponding to the 4 equality levels were included for each condition (i.e., baseline and critical), which resulted in 8 regressors of interest. By investigating GLM 1b, we can clarify how neural responses are modulated by equality signals and how such neural equality signals differ across different conditions.

In GLM 2, we tested regions computing decision utility (i.e., utility of the chosen offer) which was defined based on the winning model in the critical condition. To this end, we included the regressor corresponding to the onset of alternative offer presentation in each condition separately (i.e., baseline, critical, and filler) in the same way as GLM 1. Since we only applied computational modelling analyses in the critical condition, we only included the trial-wise utility of the chosen offer and the utility of the unchosen offer as the parametric modulators for the alternative offer events in the critical condition. The durations for these events were equal to the time form onsets of the alternative offer presentation to the time points of offsets.

To further investigate neural sensitivity to equality signals and the relationship between striatum and other brain regions modulated by equality signals, we constructed GLM 3 in which trials were divided into 2 levels of equality difference between alternative offers (i.e. high equality difference: − Δ *F* = −2 and −4, vs low equality difference: Δ *F* = −6 and −8) and 2 different alternative offer onset regressors corresponding to the 2 equality levels were included for each condition (i.e., baseline and critical). In line with GLM 1 results, GLM 3 also suggested that in the baseline condition, activity in striatum was stronger for higher equality difference than lower equality difference, and the effect was not observed in the critical. We also performed functional connectivity analyses based on GLM 3 to examine how the striatum connects with other brain regions depending on different equality levels and different conditions.

In GLM 4, we identified neural activitiy associated with specific choices during wealth redistribution in both conditions. Thus, GLM 4 included onsets of alternative offer presentation of each condition with respect to specific choice (i.e., equal choice or unequal choice in the baseline and critical condition), resulting in four regressors of interest. To control for any potential effect of utility of each offer on trials with regard to specific choice in the critical condition, the trial-wise utility of the chosen and the unchosen option were included as parametric modulators for each choice onset regressor. We also constructed GLM 4a in which the utility of the chosen and unchosen option were not included as parametric modulators. For GLM 4 modelling responses in both conditions, we excluded 11 participants who always chose one type of choice in either condition; and for analyses only involved in the critical condition, we excluded 7 participants who always chose one type of choice in the critical condition.

For the GLMs above, all parametric regressors were z-standardized before being entered into the GLM analyses. We switched off orthogonalization during model estimation to allow the parametric modulators to compete for variance. For all the GLMs, we incorporated onsets of fixation, initial offer presentation, and alternative offer presentation of trials with no response as regressors of no interest. For GLM 1, GLM 1a, and GLM 2, trail-wise difference in initial payoff between the two parties (Δ *Initial endowment*) was included as the parametric modulator for the initial offer event onset. In addition, we included six rigid body parameters as regressors of no interest, to account for head motion artifacts. Regressors of interest and no interest were convolved with a canonical hemodynamics response function (HRF).

We fed individual-level contrasts into group-level random-effect analyses with one-sample t tests to assess the neural parametric effects of signals for inequality, harm to others, and decision utility, or to compare parametric contrasts between the baseline and critical condition. Flexible factorial analyses were used to examine potential interaction effects. Correlation analyses were used to explore potential relationship between different prosocial motives (i.e., α, *β*, and δ) and neural activities. Since the distributions of α and *β* was positively skewed (skewness(α) = 0.49, and skewness(*β*) = 1.64), we normalized these two parameters by taking the square root, which was more normally distributed (skewness (α − normalized) = - 0.11, and skewness (*β* − normalized) = 0.25), as the measures of inequality aversion and harm aversion in correlation analyses. For significant results in these GLM analyses, we adopted a whole-brain corrected threshold [i.e., a combined threshold of voxel-level *p* < 0.001 uncorrected and cluster-level *p* < 0.05 family-wise error (FWE) correction] unless otherwise noted. The “Striatum” mask for ROI analyses was defined based on meta-analytic functional coactivation map of “Striatum” in the Neurosynth database (https://neurosynth.org/), and the peak MNI coordinates of Striatum [−12, 10, −6] were derived from this activation map. For GLM 1 and 5, small volume correction (SVC) was performed by using a cluster-level threshold *p* < 0.05 (FWE-corrected) and voxel-level threshold of *p* < 0.005 (uncorrected), and the small volume was defined as a sphere with 8 mm radius centered on the striatum peak MNI coordinates. For other GLMs, small volume corrections were performed by using a cluster-level threshold *p* < 0.05 (FWE-corrected) and voxel-level threshold of *p* < 0.001 (uncorrected), and the small volume was defined as a sphere with 10 mm radius centered on the striatum peak MNI coordinates.

### Functional connectivity analyses

By performing functional connectivity analyses, we aimed to address two questions: 1) whether the neural sensitivity to equality in striatum in the critical condition was suppressed by other cognitive processes, especially when harm aversion/rank reversal aversion conflict with inequality aversion; 2) whether different motives (i.e., harm aversion and rank reversal aversion) interact with striatum through different systems or interact with other neural pathways to affect redistribution decisions. To address these questions, we established two psychophysiological interaction analysis (PPI) models. To answer the first question, we took the striatum (i.e., a 6-mm radius sphere region centered at the peak MNI coordinates of [−12, 10, −6] of the meta-analytical “striatum” mask from Neurosynth) as the seed region in the PPI analyses. We conducted a PPI analysis for each of the two conditions (i.e., Baseline and Critical) to assess differential functional connectivity with this seed in high − Δ *F* trials compared with low − *Δ F* trials based on GLM 3. For each PPI analysis, the BOLD signal within the seed (i.e., average time series within 6-mm sphere around the peak voxel) was used as the physiological factor and high equality signal (− Δ *F*) versus low equality signals (− Δ *F*) contrast in GLM 3 was used as the psychological factor. At the first level, the PPI model included one regressor representing the extracted time series in the seed (i.e., the physiological variable), one regressor representing the psychological variable of interest, and a third regressor representing the interaction of the two regressors (the PPI term).

To answer the second question, we took the striatum region that was associated with equality processing and equal choice (MNI coordinates: [−18, 11, −2]) identified in GLM 1 as the seed region. Since for these PPI analyses, we aimed to examine the neural network underlying redistribution decisions, we focused on the contrast of “more unequal choice > more equal choice” in GLM 4. Therefore, the PPI models included one regressor representing the extracted time series in the seed (a 6-mm sphere region centered at coordinates corresponding to each region of striatum) as the physiological variable, one regressor representing the psychological variable of interest (i.e., more unequal choice > more equal choice), and a third regressor representing the interaction of the two regressors (the PPI term).

At the second level, for the first PPI analyses, two beta maps of the PPI term in the baseline and critical conditions for each participant were fed into a paired t-test analyses. For the second PPI analyses, since we were interested in how different motives modulate the neural networks to affect decisions, we correlated the beta maps with individuals’ parameters of inequality aversion (α), harm aversion (*β*), or rank reversal aversion (δ) derived from the winning computational model. Significant results were reported with a whole-brain corrected threshold [i.e., a combined threshold of voxel-level *p* < 0.001 uncorrected and cluster-level *p* < 0.05 family-wise error (FWE) correction] unless a special note.

## Acknowledgement

This study was supported by grants from the National Natural Science Foundation of China (31630034, 71942001). Dr. Jie Hu and Dr. Christian C. Ruff also received funding from the European Research Council (ERC) under the European Union’s Horizon 2020 research and innovation programme (grant agreement No 725355).

