## supplementary information for "Neurocomputational evidence that conflicting prosocial motives guide distributive justice"

**Supplementary methods**

**Model construction**

To formalize different motives underlying redistribution behaviors, we performed model-based analyses by developing and comparing four families of computational models to identify how people weigh between multiple motives to make redistributive decisions. It is noteworthy that we only performed computational modelling analyses for the critical condition. This is because, first, to match with the trials in the critical condition, we generated alternative offers in the baseline condition with the same inequality levels as the counterpart offers in the critical condition and there is no variance in efficiency across alternative offers since the initial offer and both alternative offers have the same sum of payoffs. Therefore, computational models given the trial set in the baseline condition could not effectively measure different levels of inequality aversion across participants. Second, as the difference in inequality and the difference in transferred money between alternative offers were completely correlated with each other in the baseline condition, it is impossible to differentiate inequality aversion from harm aversion in the baseline condition.

Then, how might participants weigh between the more equal and more unequal alterative offers in the critical condition? On possibility is that, following the classical inequality aversion model proposed by Fehr and Schmidt (1999), people will value the outcomes of all parties and devalue the inequality they experience for any kinds of distribution (Model M1). Since both parties of the distribution are anonymous strangers for the participant, we considered the absolute value of the payoff difference between the two parties as the inequality level of each offer. And, people will devalue the utility of the offer more strongly for higher inequality level. Therefore, the utilities of the two alternative offers and utility difference between the two offers ($\Delta U$) are as follows:

$$U(Unequal) = I_A + I_B - \alpha\,|I_A - I_B|$$

$$U(Equal) = E_A + E_B - \alpha\,|E_A - E_B|$$

$$\Delta U = U(Equal) - U(Unequal) = [E_A + E_B - \alpha\,|E_A - E_B|] - [I_A + I_B - \alpha\,|I_A - I_B|]$$

$$= \alpha\ (|I_A - I_B| - |E_A - E_B|)$$

where $I_A (I_B)$ is the payoff of the more unequal alternative offer for initially advantaged (disadvantaged) party, $E_A (E_B)$ is the payoff of the more equal alternative offer for initially advantaged(disadvantaged) party, and $\alpha$ is the inequality aversion parameter that captures the weighing of inequality level of the offers. Since in the current paradigm, the two alternative offers have the same payoff sum (e.g., $I_A + I_B = E_A + E_B$), the utility difference ($\Delta U$) is mainly driven by the difference in inequality level between the two offers (i.e., $|I_A - I_B| - |E_A - E_B|$). We referred this inequality difference as $\Delta F$ ($\Delta F = |I_A - I_B| - |E_A - E_B|$). Therefore,

$$\Delta U = \alpha \Delta F \qquad \text{(M1)}$$

Another possibility is that, since only the more equal alternative offer will reverse the relative rankings between the two parties in the initial offer, participants who are averse to rank reversal will devalue the utility of the more equal offer ($U(Equal)$) for rank reversal. Therefore, we included one discounting parameter $\delta$ to capture the rank reversal aversion for the more equal offer:

$$U(Equal) = E_A + E_B - \alpha\ |E_A - E_B| - \delta$$

Then,

$$\Delta U = U(Equal) - U(Unequal) = [E_A + E_B - \alpha\ |E_A - E_B| - \delta] - [I_A + I_B - \alpha\ |I_A - I_B|]$$

$$= \alpha\ (|I_A - I_B| - |E_A - E_B|) - \delta$$

$$= \alpha \Delta F - \delta \qquad \text{(M2)}$$

A third possibility is that people may also be averse to benefit one party by harming the other one. Therefore, it is plausible that the more money the alternative offer takes away from the initially advantaged party, the more averse the participant is to the offer. In other words, people will not only devalue the offers for the inequality level and rank reversal but also devalue them for the extent the offers harm the initially advantaged party. To scrutinize how people weigh the harm of alternative offers to the initially advantaged party, we constructed six more models assuming three different strategies of devaluing harms.

First, in models M3a to M3c, we assumed that people would devalue the utility of the alternative offer for the amount of money transferred from initially advantaged party to the disadvantaged party. To reach to the more equal offer, participants need to transfer a larger amount of money from initially advantaged party to disadvantaged party than to reach to the more unequal offer. Therefore, we assumed that in the critical condition, in addition to the difference in inequality level ($\Delta F$) and rank reversal, participants would also consider the difference in the amount of money transferred across the two parties between the two offers ($\Delta T$):

$$U(Unequal) = \mathrm{I}_A + I_B - \alpha\,|\mathrm{I}_A - I_B| - \beta(D_A - I_A)$$

$$U(Equal) = E_A + E_B - \alpha\,|E_A - E_B| - \beta(D_A - E_A) - \delta$$

$$\Delta U = U(Equal) - U(Unequal)$$

$$= \alpha(\,|I_A - I_B| - |E_A - E_B|) - \beta((D_A - E_A) - (D_A - I_A)) - \delta$$

$$\Delta F = |I_A - I_B| - |E_A - E_B|$$

$$\Delta T = I_A - E_A$$

$$\Delta U = \alpha\Delta F - \beta\Delta T - \delta \qquad \text{(M3a)}$$

where $D_A(D_B)$ is the payoff of the initial offer for advantaged (disadvantaged) party, $\Delta F$ is the difference in inequality level between the two alternative offers as above models, and $\Delta T$ is the difference in the amount of money transferred across the two parties between the two alternative offers. $\alpha$ and $\delta$ are still the inequality aversion parameter and rank reversal aversion parameter as control models. $\beta$ is the harm aversion parameter that captures the subjective cost to take money away from the initially advantaged party.

However, it is possible that people are not averse to transfer more money as long as the transferred money can decrease the initial inequality level. Instead, they may only be averse to reach a certain equality level by transferring more money than necessary. Therefore, in models M4a to M4c, we assumed that participants would evaluate inequality difference between alternative offers ($\Delta F$) and rank reversal of the more equal offer as control models. But, as the more equal alternative offer transferred more money than the amount that can reach the same level of equality as the offer itself but not reverse the initial rankings, the model assumed that people are averse to reach a certain equality level by transferring such extra money. We considered the proportion of extra transferred money as unnecessary loss or harm for the initially advantaged party (i.e., $H$). That is to say, people will devalue the more equal offer for the proportion of the transferred money that exceeds the necessary amount of money that can both reach the same equality level and keep the initial rankings (captured by the harm aversion parameter $\beta$). Given the current paradigm, the extra proportion of transferred money equals to the difference in payoff of the more equal offer between the initially disadvantaged and advantaged party (i.e., $E_B - E_A$). Therefore, Model M4a is as follows:

$$U(Unequal) = \mathrm{I}_A + I_B - \alpha\,|\mathrm{I}_A - I_B|$$

$$U(Equal) = E_A + E_B - \alpha\,|E_A - E_B| - \beta((D_A - E_A) - (D_A - E_B)) - \delta$$

$$\Delta U = U(Equal) - U(Unequal) = \alpha(\,|I_A - I_B| - |E_A - E_B|) - \beta(E_B - E_A) - \delta$$

$$\Delta F = |I_A - I_B| - |E_A - E_B|$$

$$H = E_B - E_A$$

$$\Delta U = U(Equal) - U(Unequal) = \alpha\Delta F - \beta H - \delta \quad \text{(M4a)}$$

where $\alpha$ and $\delta$ are the inequality aversion parameter and rank reversal parameter as previous models. $\beta$ is the harm aversion parameter that captures the subjective cost of taking more money than necessary from the advantaged party (i.e., generating greater others' loss or harm), and $H$ is the proportion of the transferred money in more equal offer that exceeds the necessary amount of money that can both reach the same equality level and keep the initially relative rankings.

To summarize, in total, we established eight models which held different assumptions about how people devalue the utility alternative offers in the critical condition. We consider M1 and M2 as control models which did not consider harm aversion and rank reversal aversion (M1), or harm aversion (M2). For models M3a-M3c and M4a-M4c, models within the same set of models shared the same way of calculation of harm, but assumed different types of devaluations of harm and rank reversal, which is already described in the Materials & Methods section.

**Model simulation**

We performed model simulation analyses to test whether and how simulated choice would vary with the three parameters of interest in the winning model (i.e., inequality aversion $\alpha$, harm aversion $\beta$, and rank reversal aversion $\delta$). Similar as the parameter recovery analysis, we generated 27 datasets using all combinations of three plausible values for each parameter (α: 0.1, 0.3, 0.6; $\beta$: 0.1, 0.3, 0.6; $\delta$: 0.8, 1.1, 1.4). The temperature parameter $\lambda$ was fixed at 1.2. We simulated agent's responses with the winning model in the critical condition 50 times for each parameter combination; and, for each repeat, a small noise derived from a uniform distribution (0, 0.1) was added to each of the three the parameter values. The simulation result is shown in Figure S3.

**Supplementary results**

In the analyses of GLM 4, we showed that greater activity in DMPFC and TPJ were associated with stronger inequality aversion, and that greater activity in putamen was associated with stronger harm aversion when people choosing the more unequal offer. These correlation patterns still held after controlling for the effect of the other two parameters (for DMPFC – $\alpha$ [$\beta$ and $\delta$ controlled]: MNI peak coordinates: [ -3, 50, 34], max t-value = 3.46, *p*(FWE-SVC) = 0.022; for TPJ – $\alpha$ [$\beta$ and $\delta$ controlled]: MNI peak coordinates: [ -57, -64, 25], max t-value = 3.46, *p*(FWE-SVC) = 0.018; for Putamen – $\beta$ [$\alpha$ and $\delta$ controlled]: MNI peak coordinates: [ -18, 8, -5], max t-value = 3.56, *p*(FWE-SVC) = 0.008).

In the PPI analyses of GLM 4, we revealed association between stronger IFG-striatum connectivity and greater inequality aversion, association between stronger SFG-striatum connectivity and greater rank reversal aversion, and association between stronger STG-IPL connectivity and greater harm aversion. These correlation patterns of different networks also held after controlling for the effect of the other two parameters (for striatum-IFG with $\alpha$ [$\beta$ and $\delta$ controlled]:  MNI peak coordinates: [57, 23, 4], max t-value = 4.53, *p*(FWE-SVC) = 0.006; for striatum-SFG with $\delta$ [$\alpha$ and $\beta$ controlled]:  MNI peak coordinates: [-18, 5, 49], max t-value = 4.14, *p*(FWE-SVC) = 0.008; for IPL-STG with $\beta$ [$\alpha$ and $\delta$ controlled]: MNI peak coordinates: [ -63, -13, 10], max t-value = 4.22, *p*(FWE) = 0.018).

**Table S1.** Mixed-effects model results of behavioral data in fMRI wealth redistribution task

| | Generalized mixed-effects model | | |
|---|---|---|---|
| Variable | *B (S.E.)* | *ORE (95% CI)* | *P Value* |
| Intercept | 1.15(0.18) | 3.17(2.21, 4.53) | < 0.001 |
| Condition (C) | -0.77 (0.04) | 0.37 (0.33, 0.42) | < 0.001 |
| Δ Inequality (I) | 0.46 (0.07) | 1.58 (1.37, 1.83) | < 0.001 |
| Δ Initial Endowment (E) | 0.11 (0.05) | 1.12 (1.01, 1.24) | 0.04 |
| Δ Transfer (T) | -0.77 (0.04) | 0.46 (0.43, 0.50) | < 0.001 |
| C*I | -0.14 (0.09) | 0.87 (0.73, 1.03) | 0.11 |
| C*E | -0.02 (0.05) | 0.98 (0.88, 1.09) | 0.70 |
| I*E | -0.06 (0.07) | 0.94 (0.82, 1.09) | 0.44 |
| I*T | -0.37 (0.16) | 0.69 (0.50, 0.96) | 0.03 |
| E*T | 0.17 (0.04) | 1.19 (1.10, 1.28) | < 0.001 |
| C*I*E | -0.02 (0.09) | 0.98 (0.82, 1.16) | 0.79 |
| C*I*T | 0.37 (0.11) | 1.44 (1.16, 1.79) | < 0.001 |
| I*E*T | 0.11(0.16) | 1.12 (0.82, 1.53) | 0.48 |
| C*I*E*T | -0.11 (0.11) | 0.90 (0.72, 1.11) | 0.31 |
| LL | | -4642 | |
| BIC | | 9422 | |
| Marginal $R^2$ | | 0.19 | |

ORE, odds ratio estimate; CI, confidence interval; LL, log-likelihood; BIC, Bayesian Information Criterion

**Table S2.** Main effect of Δ Inequality on probability to choose more equal alternative offer

| Δ Inequality | P (Equal choice) (Mean ± SE) |
|---|---|
| 2 | 0.52 ± 0.02 |
| 4 | 0.58 ± 0.03 |
| 6 | 0.62 ± 0.03 |
| 8 | 0.61 ± 0.03 |

**Table S4.** Main effect of Δ Transfer on probability to choose more equal alternative offer

| Δ Transfer | P (Equal choice) (Mean ± SE) |
|---|---|
| 1 - 3 | 0.76 ± 0.03 |
| 4 - 6 | 0.522 ± 0.03 |
| 7 - 11 | 0.36 ± 0.04 |

**Table S5.** Interaction between Δ Inequality and Δ Transfer on probability to choose more equal alternative offer in the critical condition

| | Δ Transfer | | |
|---|---|---|---|
| Δ Inequality | Low (3-5) (Mean ± SE) | Middle (6-8) (Mean ± SE) | High (9-11) (Mean ± SE) |
| 2 | 0.27 ± 0.04 | 0.28 ± 0.05 | 0.27 ± 0.04 |
| 4 | 0.38 ± 0.05 | 0.39 ± 0.05 | 0.36 ± 0.05 |
| 6 | 0.46± 0.05 | 0.42 ± 0.05 | 0.45 ± 0.05 |
| 8 | 0.51 ± 0.05 | 0.40 ± 0.05 | 0.41 ± 0.05 |

**Table S6.** Interaction between Δ Initial Endowment and Δ Transfer on probability to choose more equal alternative offer in the critical condition

| | Δ Transfer | | |
|---|---|---|---|
| Δ Initial Endowment | Low (3-5) (Mean ± SE) | Middle (6-8) (Mean ± SE) | High (9-11) (Mean ± SE) |
| Low | 0.76 ± 0.03 | 0.49 ± 0.03 | 0.34 ± 0.04 |
| High | 0.75 ± 0.02 | 0.59 ± 0.03 | 0.38 ± 0.04 |

Table S7. Quality of model fits for computational models of redistribution decision-making.

| **Model** | **Description** | **Parameters per subject** | **BIC** | **Cross-validated prediction accuracy (Mean ± SE)** |
|---|---|---|---|---|
| M1 | $\alpha, \lambda$ | 2 | 4417 | 0.478 ± 0.025 |
| M2 | $\alpha, \delta, \lambda$ | 3 | 3458 | 0.536 ± 0.021 |
| M3a | $\alpha, \beta, \delta, \lambda$ | 4 | 2677 | 0.744 ± 0.025 |
| M3b | $\alpha, \beta, \lambda$ | 3 | 2931 | 0.726 ± 0.025 |
| M3c | $\alpha, \beta, \delta, \lambda$ | 4 | 2811 | 0.738 ± 0.024 |
| M4a | $\alpha, \beta, \delta, \lambda$ | 4 | 2639 | 0.749 ± 0.025 |
| M4b | $\alpha, \beta, \lambda$ | 3 | 2869 | 0.726 ± 0.025 |
| M4c | $\alpha, \beta, \delta, \lambda$ | 4 | 2808 | 0.734 ± 0.024 |

BIC, Bayesian Information Criterion. Model M4a was favored. All models have inverse temperature parameter $\lambda$. $\alpha$, Inequality aversion parameter; $\beta$, Harm aversion parameter; $\delta$, Rank reversal aversion parameter.

**Table S8.** Results of whole-brain/ROI parametric analysis of fMRI data in GLM 1

| Regions | Laterality | Peak MNI coordinates | | | Max T-value | Cluster size (k) |
|---|---|---|---|---|---|---|
| | | x | y | z | | |
| Baseline (whole-brain): Positive association with $-\Delta F$ | | | | | | |
| Caudate/Putamen | L | -18 | 5 | 19 | 4.24 | 290 |
| Lingual gyrus | R | 21 | -73 | -8 | 8.03 | 1521 |
| MOG | L | -27 | -97 | 4 | 4.52 | 258 |
| Critical (whole-brain): Positive association with $-\Delta F$ | | | | | | |
| No significant cluster | | | | | | |
| Baseline > Critical (whole-brain): Association with $-\Delta F$ | | | | | | |
| Lingual gyrus | R | 12 | -79 | -2 | 5.70 | 683 |
| Critical (whole-brain): Positive association with harm to others ($H$) | | | | | | |
| IPL | L | -39 | -49 | 46 | 8.21 | 3516 |
| | R | 39 | -49 | 46 | | |
| DMPFC/ACC | L | -6 | 29 | 34 | 6.90 | 657 |
| IFG | R | 48 | 17 | 31 | 6.86 | 1351 |
| MFG | L | -42 | 11 | 52 | 6.17 | 921 |
| Fusiform Gyrus | L | -36 | -58 | -14 | 5.87 | 470 |
| ITG | R | 54 | -49 | -23 | 4.96 | 292 |
| Baseline (Striatum ROI): Positive association with $-\Delta F$ | | | | | | |
| Caudate/Putamen | L | -18 | 11 | 1 | 3.64 | 111 |
| | R | 15 | 20 | -5 | 3.55 | 76 |
| Baseline > Critical (Striatum ROI): Association with $-\Delta F$ | | | | | | |
| Caudate | R | 6 | 14 | -5 | 4.01 | 45 |

MOG, middle occipital gyrus; IPL, inferior parietal lobe; DMPFC, dorsolateral prefrontal cortex; ACC, anterior cingulate cortex; IFG, inferior frontal gyrus; MFG, middle frontal gyrus; ITG, inferior temporal gyrus. ROI, regions of interest. $\Delta F$, inequality difference between alternative offers; $H$, harm amount, the proportion of the transferred money that exceeds the necessary amount of money that can reach the same equality level but not reverse the initial relative rankings for the more equal alternative offer. Significant clusters were thresholded at voxel- wise $p < 0.001$ uncorrected and cluster-wise FWE corrected $p < 0.05$.

**Table S9.** Results of whole-brain/ROI parametric analysis of fMRI data in GLM 1a

| Regions | Laterality | Peak MNI coordinates x | y | z | Max T-value | Cluster size (k) |
|---|---|---|---|---|---|---|
| Baseline (whole-brain): Positive association with $-\Delta F$ | | | | | | |
| Caudate/Putamen | L | -18 | 5 | 19 | 4.25 | 301 |
| Lingual | R | 21 | -73 | -8 | 8.11 | 1571 |
| MOG | L | -27 | -97 | 4 | 4.55 | 274 |
| Critical (whole-brain): Positive association with $-\Delta F$ | | | | | | |
| No significant cluster | | | | | | |
| Baseline (Striatum ROI): Positive association with $-\Delta F$ | | | | | | |
| Caudate/Putamen | L | -18 | 11 | 1 | 3.98 | 120 |
| | R | 15 | 20 | -5 | 3.82 | 90 |
| Baseline: $-\Delta F$ > Critical: $-\Delta F$ (Striatum ROI) | | | | | | |
| Caudate | L | -18 | 17 | 2 | 3.44 | 7 |
| | R | 6 | 14 | -5 | 3.39 | 18 |

MOG, middle occipital gyrus. $\Delta F$, inequality difference between alternative offers. Significant clusters were thresholded at voxel- wise $p < 0.001$ uncorrected and cluster-wise FWE corrected $p < 0.05$.

**Table S10.** Results of whole-brain analysis of fMRI data in GLM 4

| Regions | Laterality | Peak MNI coordinates | | | Max T-value | Cluster size (k) |
|---|---|---|---|---|---|---|
| | | x | y | z | | |
| Baseline: unequal choice > equal choice | | | | | | |
| MFG/IFG/Insula | R | 51 | 20 | 16 | 5.70 | 2586 |
| | L | -36 | 29 | 28 | 5.20 | 374 |
| TPJ | R | 51 | -46 | 31 | 4.78 | 208 |
| Baseline: equal choice > unequal choice | | | | | | |
| No significant cluster | | | | | | |
| Critical: unequal choice > equal choice | | | | | | |
| No significant cluster | | | | | | |
| Critical: equal choice > unequal choice | | | | | | |
| No significant cluster | | | | | | |
| Critical: unequal choice > equal choice, positively correlated with inequality aversion ($\alpha$) | | | | | | |
| TPJ | L | -60 | -55 | 25 | 5.58 | 266 |
| DMPFC | L | -3 | 56 | 28 | 4.14 | 161 |
| Critical: unequal choice > equal choice, positively correlated with harm aversion ($\beta$) | | | | | | |
| Putamen | L | -18 | 8 | -5 | 3.88 | 137 |
| MOG | R | 21 | -94 | 7 | 4.45 | 276 |
| IOG | L | -51 | -61 | -14 | 4.59 | 415 |

MFG, middle frontal gyrus; IFG, inferior frontal gyrus; TPJ, temporoparietal junction; DMPFC, dorsolateral prefrontal cortex; MOG, middle occipital gyrus; IOG, inferior occipital gyrus. Significant clusters were thresholded at voxel- wise $p < 0.001$ uncorrected and cluster-wise FWE corrected $p < 0.05$.

**Table S11.** Results of whole-brain analysis of fMRI data in GLM 4a

| Regions | Laterality | Peak MNI coordinates | | | Max T-value | Cluster size (k) |
|---|---|---|---|---|---|---|
| | | x | y | z | | |
| Baseline: unequal choice > equal choice | | | | | | |
| MFG/IFG | R | 51 | 20 | 16 | 5.56 | 1929 |
| | L | -36 | 29 | 28 | 5.17 | 340 |
| Insula | L | -27 | 26 | -5 | 5.20 | 605 |
| TPJ | R | 51 | -46 | 31 | 4.81 | 228 |
| IPL | L | -45 | -52 | 40 | 4.18 | 147 |
| Baseline: equal choice > unequal choice | | | | | | |
| No significant cluster | | | | | | |
| Critical: unequal choice > equal choice | | | | | | |
| No significant cluster | | | | | | |
| Critical: equal choice > unequal choice | | | | | | |
| No significant cluster | | | | | | |

MFG, middle frontal gyrus; IFG, inferior frontal gyrus; TPJ, temporoparietal junction; IPL, inferior parietal lobe. Significant clusters were thresholded at voxel- wise $p < 0.001$ uncorrected and cluster-wise FWE corrected $p < 0.05$.

**Table S12.** Results of PPI analysis of unequal choice vs equal choice in the critical condition

| Regions | Laterality | Peak MNI coordinates x | y | z | Max T-value | Cluster size (k) |
|---|---|---|---|---|---|---|
| **PPI**: unequal choice > equal choice, seed striatum centered at [-18, 11, -2] | | | | | | |
| Positively correlated with inequality aversion ( α) | | | | | | |
| IFG | R | 57 | 26 | 13 | 5.34 | 113 |
| Positively correlated with rank reversal aversion (δ) | | | | | | |
| SFG | L | -21 | -1 | 49 | 5.46 | 165 |

IFG, inferior frontal gyrus; SFG, superior frontal gyrus; STG, superior temporal gyrus. Significant clusters were thresholded at voxel- wise $p < 0.001$ uncorrected and cluster-wise FWE corrected $p < 0.05$.

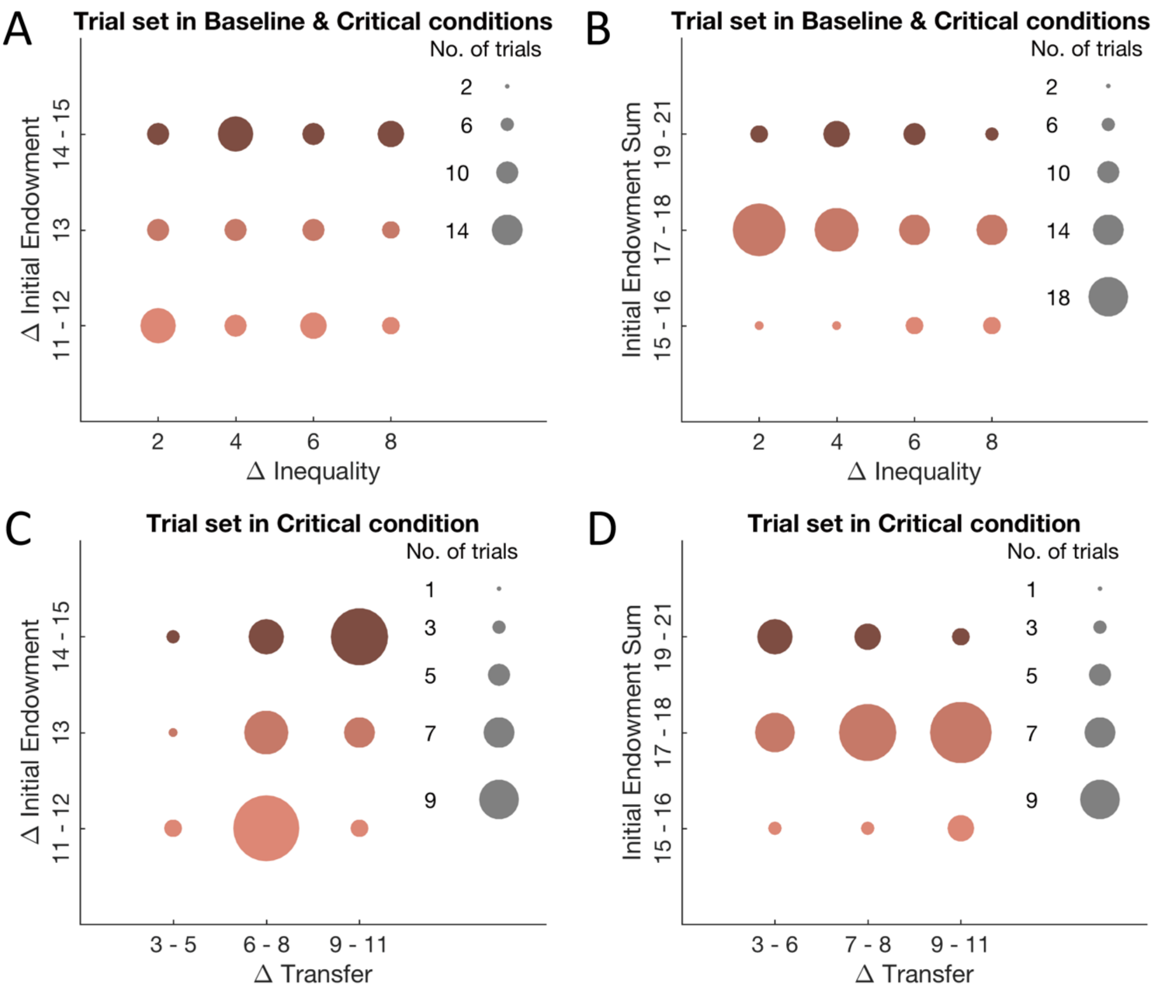


**Figure S1. Trial set matrix of the baseline and critical conditions.** To clarify the effects of inequality and the amount of transferred money from the advantaged party, we orthogonalized the differences in inequality/transferred money between the two alternative offers with the difference/sum of the initial endowment of the two parties. (A & B) Design matrix with x axis representing the difference in inequality level (Δ Inequality) between the two alternative offers, and y axis representing the difference between the two parties in initial endowment (A) and the sum of initial endowment for the two parties (B). (C & D) Design matrix with x axis representing the difference in transferred money (Δ Transfer) between the two alternative offers, and y axis representing the difference between the two parties in initial endowment (C) and the sum of initial endowment for the two parties (D). The size of the circle is proportional to the number of trials in each type of variable combination.

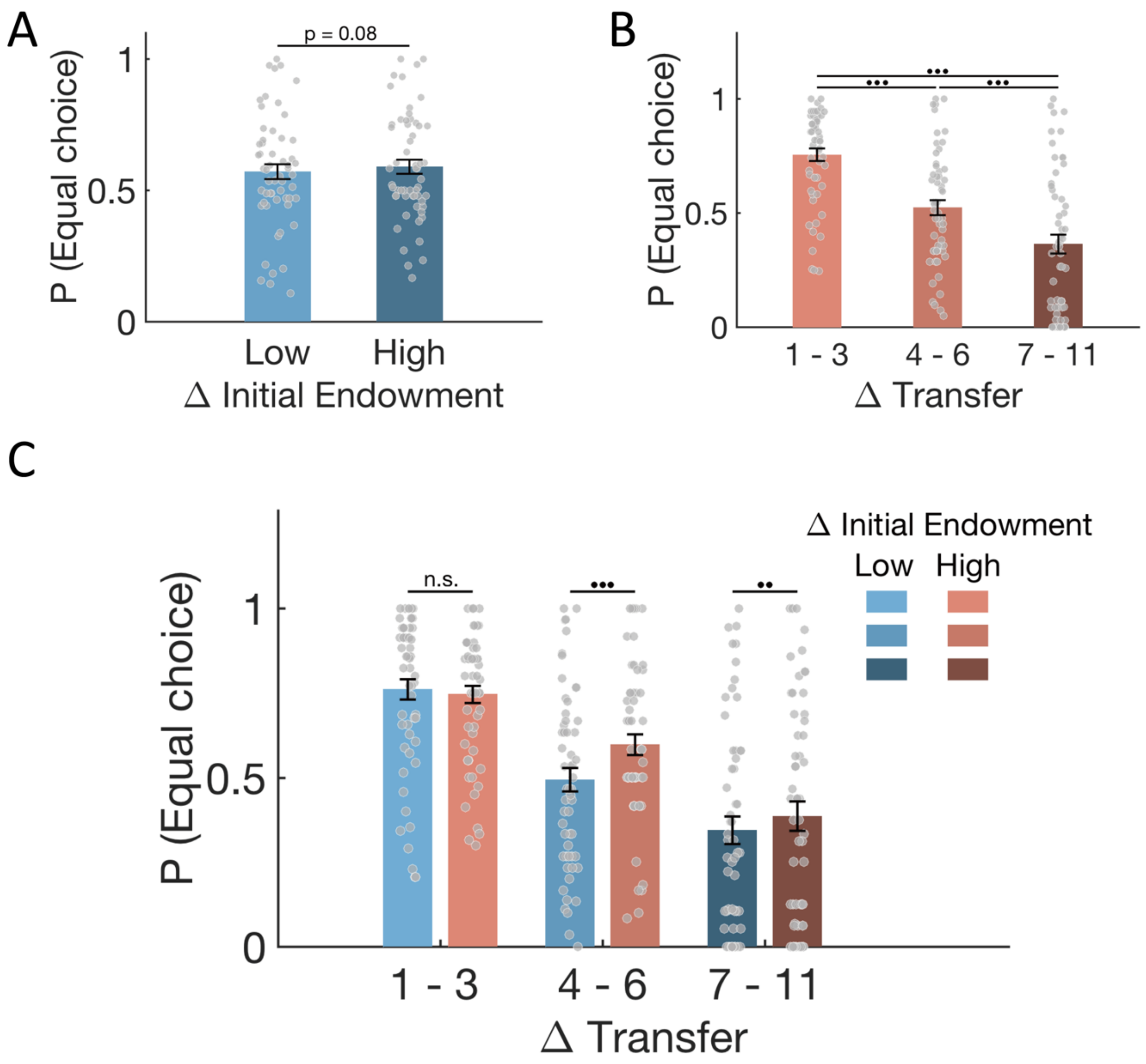


**Figure S2. Behavioral results. (A)** Main effect of difference in initial endowment (∆ Initial endowment) between the two parties on probability to choose more equal offer. Participants chose more equal offer more frequently when initial endowment difference was higher. **(B)** Main effect of difference in transferred money (∆ Transfer) between the two alternative offers on probability to choose more equal offer. Probability to choose more equal offer decreased with the increase of difference in transferred money between the more equal offer and the more unequal offer. **(C)** Interaction between ∆ Initial endowment and ∆ Transfer on probability to choose more equal offer. When the difference in transferred money is high (i.e., 4 - 6 and 7 - 11), higher initial endowment difference increased the probability to choose more equal offer. Each grey dot represents one participant, and error bars represent the SEMs. •••, $p < 0.001$; ••, $p < 0.01$; •, $p < 0.05$, n.s., $p > 0.1$.

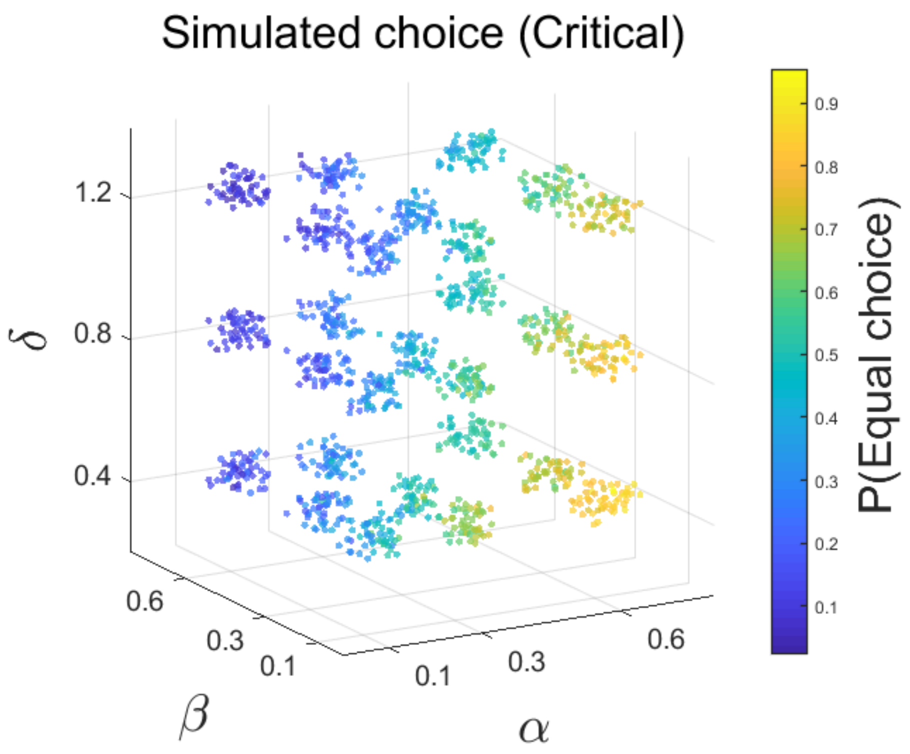


**Figure S3. Simulation results of the winning model.** The 3-D scatter plot shows that the simulated responses [i.e., P(Equal choice)] vary with the three parameters, indicating that different motives affect individuals' redistribution decisions in different manners. To do this analysis, we generated 27 datasets using all combinations of three plausible values for each parameter ($\alpha$: 0.1, 0.3, 0.6; $\beta$: 0.1, 0.3, 0.6; $\delta$: 0.8, 1.1, 1.4). The temperature parameter $\lambda$ was fixed at 1.2. Each dot represents one repeat of simulation. For each repeat, a small noise derived from a uniform distribution (0, 0.1) was added to each of the three the parameter values in a certain parameter combination. We simulated responses with the winning model in the critical condition 50 times for each parameter combination. The color of each point indicates the probability of more equal choice in each simulation.

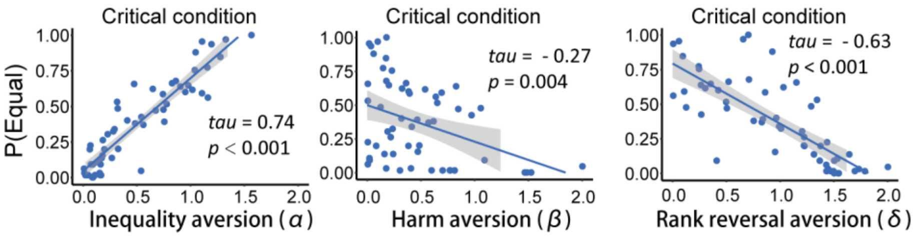


**Figure S4. Correlations between probability to choose the more equal offer and model parameters which correspond to different motives in the critical condition.** Scatter plots show that individuals who are more averse to inequality (i.e., higher α, left panel) will choose the more equal offer more frequently, and individuals who are more averse to harming others (i.e., higher $\beta$, middle panel) and rank reversal (i.e., higher δ, right panel) will choose the more equal offer less frequently. Each dot represents one participant.

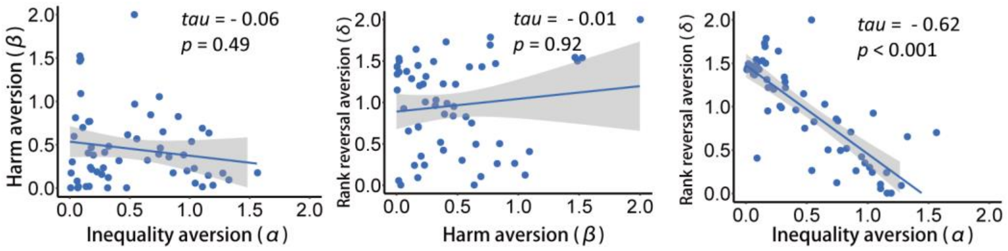


**Figure S5. Correlation analyses between the three model parameters in the winning model (M4a).** Scatter plots show that α (inequality aversion) and δ (rank reversal aversion) are negatively associated with each other. Each dot represents one participant.

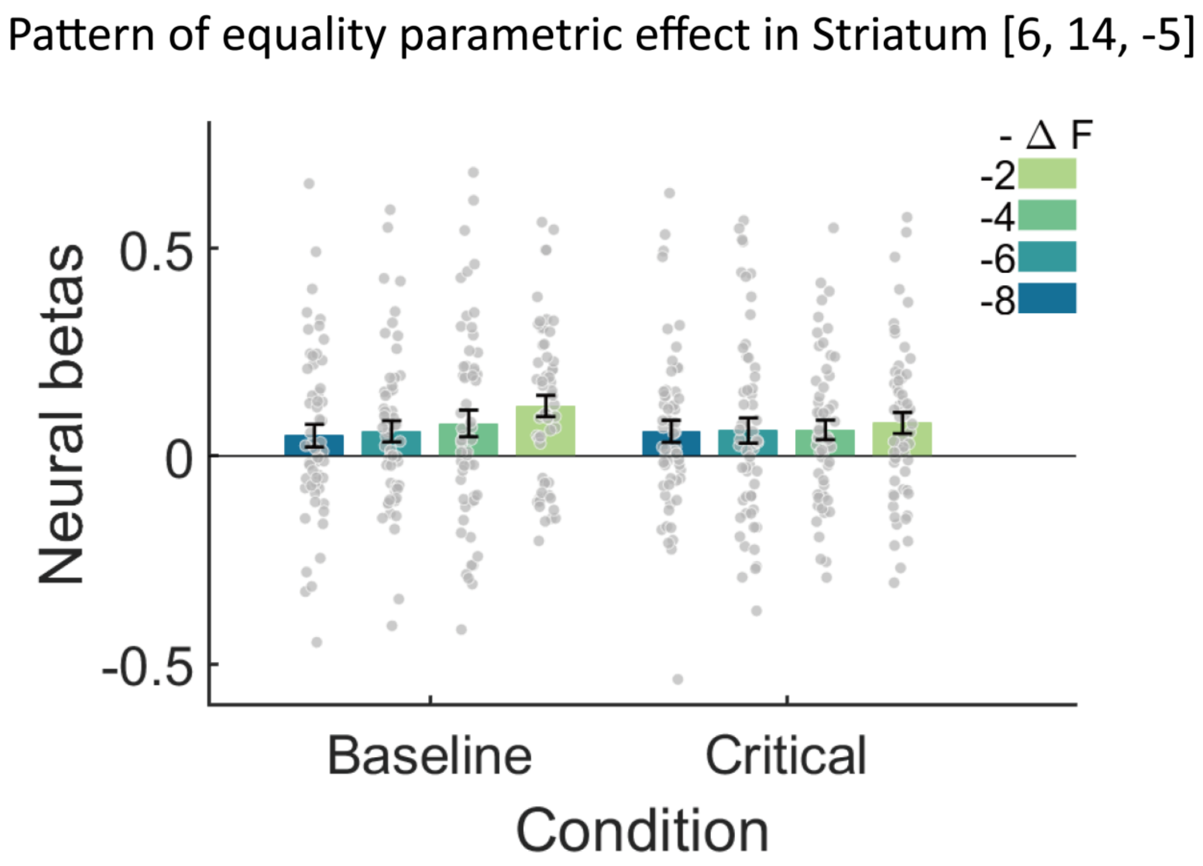


**Figure S6. Response pattern for parametric effects of equality in striatum identified in GLM 1.** To visualize the effect of parametric modulation of inequality difference identified in GLM 1, we separately regressed the four levels of equality difference (i.e., -ΔF = -2, -4, -6, and -8) in each condition, and constructed and re-estimated GLM 1b and extracted neural betas in the significant cluster (i.e., caudate/putamen) identified in GLM 1. In line with GLM 1 results, activity in striatum increased with the increase of equality signal (-ΔF) in the baseline condition, but no such effect was observed in the critical condition. To avoid double dipping the data, we did not perform any statistical analyses here. Each dot represents one participant, and error bars represent the SEMs.

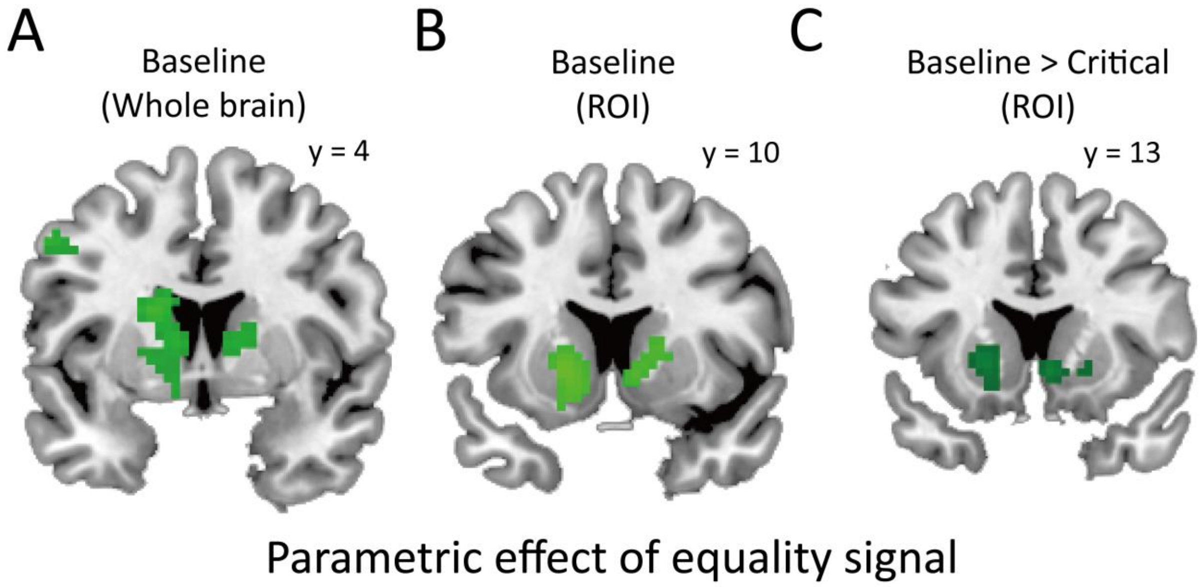


**Figure S7. Parametric analyses results in GLM 1a**. Activity in caudate/putamen was associated with the equality signal (-ΔF) in the baseline condition for both whole-brain **(A)** and independent ROI **(B)** analyses. **(C)** ROI analyses further show a greater parametric strength of equality in striatum in the baseline condition than in the critical condition. Results of GLM 1a show similar results as GLM 1, confirming the parametric effects of ΔF identified in GLM 1.

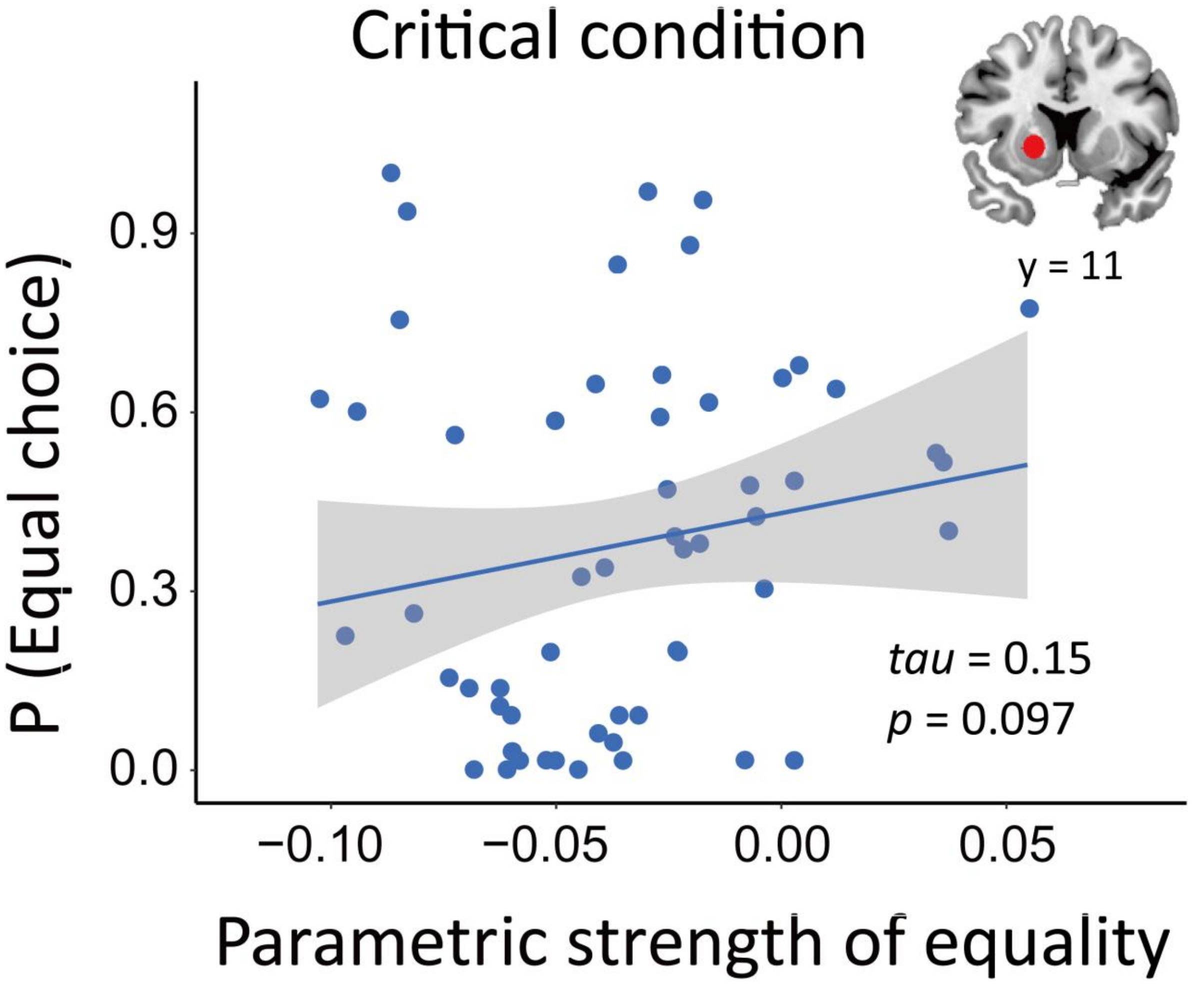


**Figure S8. Correlation between the parametric strength of equality in striatum and the probability to choose more equal offer in the critical condition.** Scatter plot shows that the correlation between the parametric strength of equality in striatum (a region with the center of peak MNI coordinates: [-12, 10, -6] in the "striatum" mask from Neurosynth) and individuals' probability to choose more equal offer in the critical condition is not significant. Each dot represents one participant.

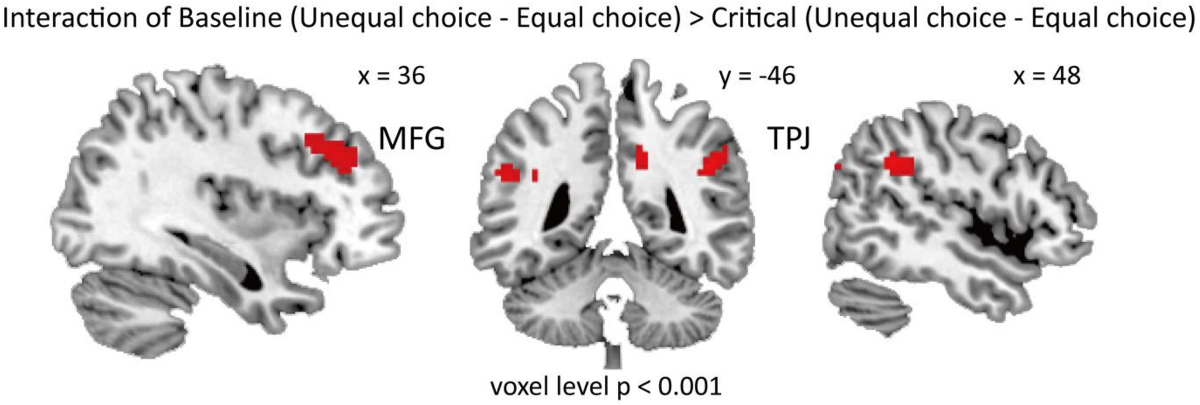


**Figure S9. Interaction between condition and choice**. A flexible factorial analysis of the interaction between condition and choice suggests that activity in MFG and TPJ are enhanced when people choosing more unequal offer than choosing more equal offer in the baseline condition, but not in the critical condition. Significant clusters were threholded with small volume correction $p$(FWE) < 0.05 with voxel-level $p < 0.001$. For purpose of visualization, results are thresholed with voxel-level $p$(uncorrected) < 0.001.

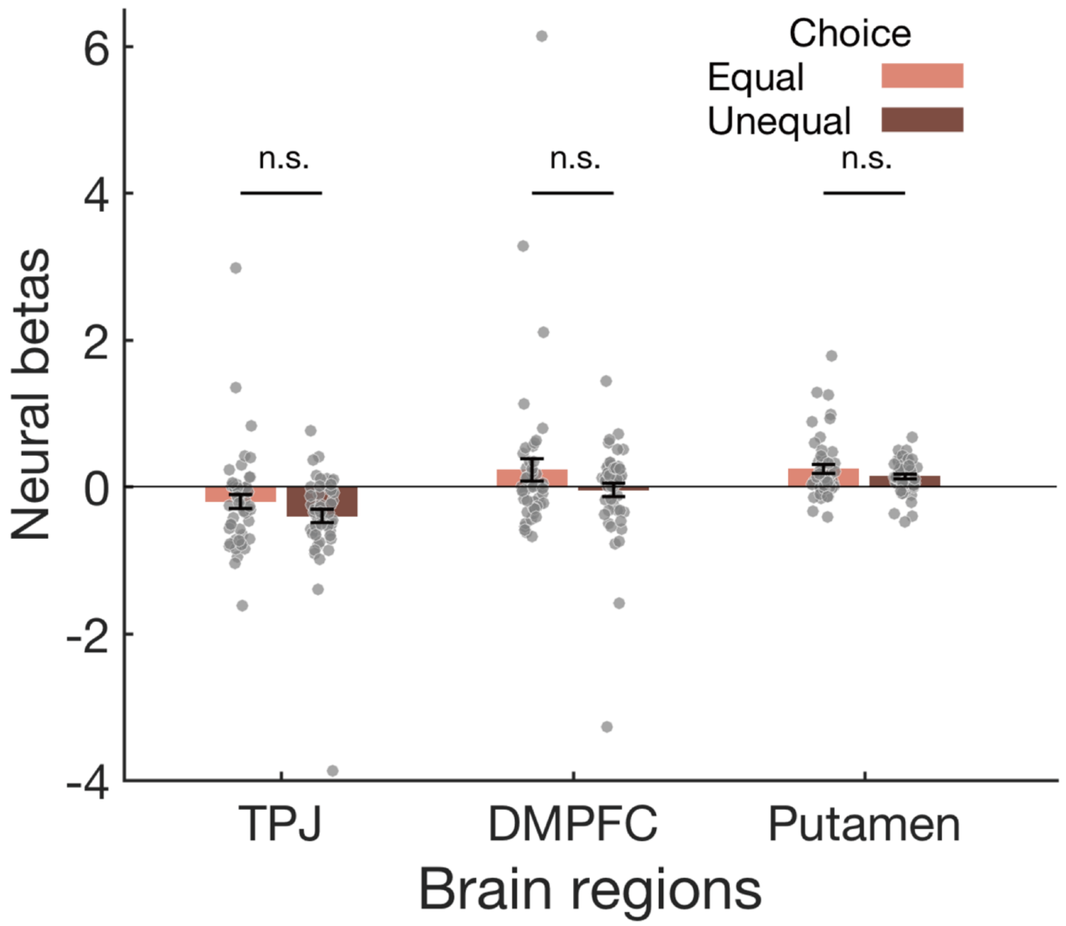


**Figure S10. No group-level difference in neural responses between choosing unequal offer and choosing equal offer in the critical condition**. For regions whose activity were shown to be associated with model parameters (i.e., $\alpha$ and $\beta$) in the contrast of "Critical: unequal choice > equal choice", there was no significant difference in their responses between choosing unequal offer and choosing equal offer in the critical condition at group level. Each dot represents one participant, and error bars represent the SEMs. n.s., not significant.
